# Maize *COI1* quadruple-knockout mutants exhibit elevated DELLA protein accumulation, stunted growth, and reduced photosynthetic efficiency

**DOI:** 10.1101/2023.04.21.537853

**Authors:** Leila Feiz, Christine Shyu, Shan Wu, Kevin R. Ahern, Iram Gull, Ying Rong, Caroline J. Artymowicz, Miguel A. Piñeros, Zhangjun Fei, Thomas P. Brutnell, Georg Jander

## Abstract

The F-box protein Coronatine Insensitive (COI) is a receptor for the jasmonic acid signaling pathway in plants. To investigate the functions of the six maize COI proteins (COI1a, COI1b, COI1c, COI1d, COI2a, and COI2b), we made single, double, and quadruple loss-of-function mutants. Double-mutant *coi2a coi2b* pollen was inviable, and no homozygous mutant plants were obtained. The *coi1* quadruple mutant (*coi1-4x*) exhibited shortened internode lengths, decreased photosynthesis, leaf discoloration, microelement deficiencies, and accumulation of DWARF9, a DELLA-family protein that represses the gibberellic acid signaling pathway. Co-expression of maize *COI* and *DWARF9* genes in *Nicotiana benthamiana* showed that the COI proteins lead to proteasome-dependent DELLA degradation. Many genes expressed at lower levels in the *coi1-4x* mutant are normally induced by gibberellic acid. The majority of these genes are predicted to be bundle sheath or mesophyll-enriched including those encoding C_4_-specific photosynthetic enzymes. Ectopic expression of maize *COI* genes in *N. benthamiana* showed that COI2a is fully localized in the nucleus and interacts with maize JAZ proteins, the canonical COI repressor partners. However, maize COI1a and COI1c proteins showed only partial nuclear localization and failed to bind to most of the JAZ proteins tested. These results show divergent functions of the six COI proteins in the regulation of maize growth and defense pathways.

## Introduction

Jasmonic acid (JA) is a lipid-derived plant hormone that regulates a wide range of biological processes, including reproductive development, vegetative growth, and responses to biotic and abiotic stresses (Qi et al., 2022; Feys et al., 1994; He et al., 2021; McConn et al., 1997; Shikha et al., 2022; Stintzi and Browse, 2000; Vijayan et al., 1998; Wang et al., 2020). Methyl JA (MeJA) is a volatile compound readily taken up and converted to JA by plants (Chuang et al., 2014). In *Arabidopsis thaliana* (Arabidopsis), *Solanum lycopersicum* (tomato), and other plants, JA is conjugated to isoleucine to produce JA-isoleucine (JA-Ile). This compound is perceived by a receptor, Coronatine Insensitive1 (COI1) (Xie et al., 1998; Xu et al., 2002), which is the F-box domain protein component of an E3-ligase complex that polyubiquitinylates JAZ repressors (Thines et al., 2007; Chini et al., 2007; Sheard et al., 2010). JAZ is a large family of ZIM-domain-containing proteins that interact with MYC transcription factors (Zhang et al., 2015; Zander et al., 2020) and thereby prevent the induction of a cohort of defense, stress, and development-related genes. Binding of JA-Ile to the COI1 receptor promotes COI1-JAZ protein-protein interactions, which in turn leads to the formation of the E3 ubiquitin ligase SCF^COI1^ complex and the subsequent degradation of the JAZ by the 26S proteasome (Fonseca et al., 2009). The de-repression of MYC transcription factors by JAZ degradation initiates transcription of the targeted genes (Dombrecht et al., 2007).

Experiments with several plant species show that JA-Ile perception and the resulting induced defense responses antagonistically modulate gibberellic acid (GA)-mediated growth (Machado et al., 2017; Campos et al., 2016; Yang et al., 2012). This reciprocal relationship between defense and growth is not due to a carbohydrate-resource trade-off (Campos et al., 2016; Machado et al., 2017) but instead results from the fact that JAZ repressors of the JA-responsive genes bind to and entrap DELLA, which represses GA-responsive growth genes, thereby preventing its repressive effect on the growth-promoting PIF transcription factors (Hou et al., 2010; Yang et al., 2012). In Arabidopsis, JA-Ile perception by COI leads to JAZ degradation, which releases DELLA and inhibits growth while inducing the transcription of defense genes by MYC transcription factors. Interruption of JA signaling in both the Arabidopsis *coi* and the rice *coi1a coi1b* mutants results in a continuous DELLA entrapment by JAZ and a positive growth response. DELLA also is essential for enhancing JA-mediated responses in Arabidopsis (Wild et al., 2012; de Vleesschauwer et al., 2016). The constitutively active, dominant DELLA mutant *gai* is hypersensitive to JA-gene induction. A quadruple *della* mutant (which lacks four of the five Arabidopsis DELLA proteins) was partially insensitive to gene induction by JA (Navarro et al., 2008).

Maize (*Zea mays*) is one of the main staple crops and a model monocot. However, both the mechanisms of JA signaling and the functions of individual maize COI receptors remain under-investigated. Whereas Arabidopsis and tomato have only one *COI* gene, gene duplication events led to the evolution of three genes in Poaceae (grasses) (Figure 1A). Modern maize has six *COI* genes (*COI1a*, *COI1b*, *COI1c*, *COI1d*, *COI2a*, and *COI2b*; Table S1), which may be due to a tetraploidization event in an ancestor of modern maize, which duplicated these genes. This tetraploidization event, which occurred between 8.5 and 13 million years ago, was followed by genome reduction of the allotetraploid maize to a diploid state with uneven gene losses and residual duplication of many genes, including the JA synthesizing enzymes (reviewed in (Borrego and Kolomiets, 2016)). CRISPR/Cas9-mediated knockouts of *COI2a* and *COI2b* showed that these genes are essential for jasmonate signaling and pollen formation in maize (Qi et al., 2022). When transformed into an *A. thaliana coi1-1* mutant, maize *COI1a*, *COI1b*, and *COI1c* but not *COI2a*, complemented the defects in fertility and induced changes in in gene expression after JA treatment (An et al., 2019). Maize COI1a interacts with JAZ15, and *coi1a* mutant plants showed elevated resistance to *Fusarium graminearum* (Gibberella stem rot) (Ma et al., 2021).

**Figure 1.**
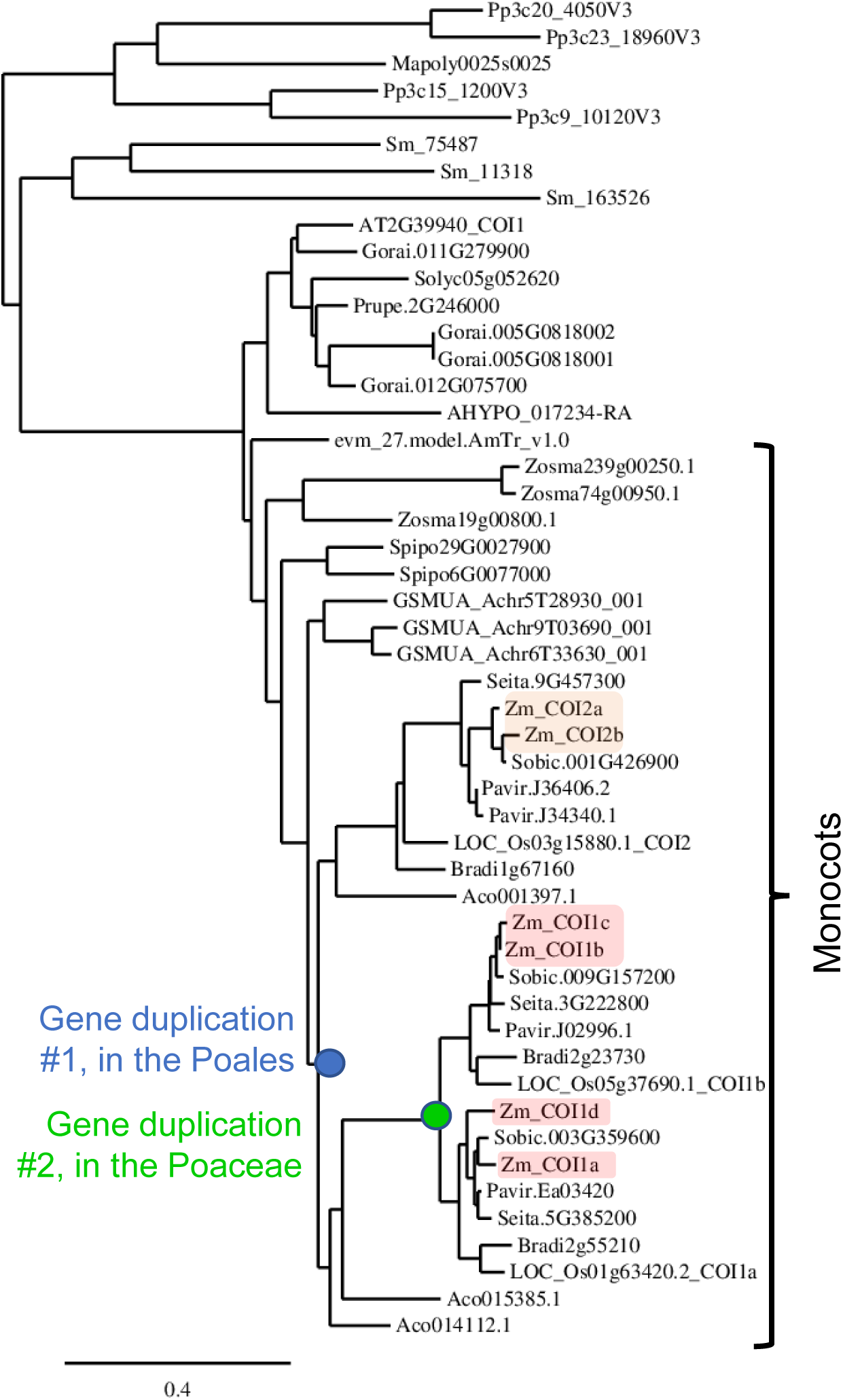
The maize genome encodes six COI proteins. Phylogenetic tree of COI proteins from maize and other species. The sequences of six maize COI proteins, along with their paralogs from other organisms, were obtained from Phytozome (https://phytozome-next.jgi.doe.gov/). The protein sequences and full species names are presented in Table S2.

Rice has three *COI* genes and, similar to maize, rice COI2 has essential roles in pollen vitality and male fertility (Inagaki et al., 2022; Trang Nguyen et al., 2022; Wang et al., 2023; Yang et al., 2012). One study showed that rice COI2 is the essential receptor in JA-mediated signaling, whereas COI1a and COI1b play redundant roles with COI2 in plant growth regulation. (Inagaki et al., 2022). In another study, rice COI2, COI1a and COI1b were all essential for JA-mediated defense against *Nilaparvata lugens*, the brown planthopper (Wang et al., 2023). In both of these studies, the rice COI2 protein showed interaction with more members of the rice *JAZ* gene family than COI1.

Here, we present a functional analysis of the maize COI proteins using transposon mutagenesis. *Ds* and *Mu* transposon insertion mutations in each of the six *COI* genes were isolated and single and higher order mutants characterized through insect bioassays, measurement of morphological changes, quantification of gene expression profiles, subcellular localization, and protein-protein interaction studies. Together, our results show that, whereas the two maize COI2 proteins have a function similar to what has been observed for Arabidopsis COI1, the four maize COI1 proteins function primarily in regulating GA-mediated growth responses. This reveals a novel molecular model where subfunctionalization of COI gene family members in maize enables growth responses to be uncoupled from plant defense pathways.

## Results

### *COI* gene duplication in monocots and maize

We identified six paralogous *COI* genes in maize, which are derived from gene duplication events in the course of monocot evolution (Figure 1; protein sequences in Table S2). These six proteins share 55% to 58% sequence identity with Arabidopsis COI1 (Figure S1A). The two members of each protein pair, COI1a and COI1d, COI1b and COI1c, and COI2a and COI2b, respectively, shared 94%, 93%, and 95% amino acid sequence identity, respectively (Figure S1B). COI1a and COI1d showed about 80% identity with COI1c and COI1b, whereas the four COI1 proteins only share about 60% identical amino acids with the two COI2 proteins. An alignment of COI protein sequences shows that phenylalanine 89 of Arabidopsis COI, which is essential for the insertion of JA-Ile between COI and JAZ (Sheard et al., 2010) is modified to tyrosine in the COI1A and COI1B clades across the Poaceae (the first red arrow in Figure S2). The only other active site amino acid changes in maize COI proteins relative to Arabidopsis are valine 411 to isoleucine in the COI2 proteins and alanine in the COI1 proteins. The other amino acids in the active site of Arabidopsis COI were not different from their corresponding amino acids in the Poaceae COI proteins (Figure S2, black arrows). Many other amino acids that are not located in the active site differ between Arabidopsis and maize COI proteins. However, based on the amino acid sequence alone, we cannot predict the function of each individual COI protein in the Poaceae.

### COI sub-cellular localization and JAZ protein interactions

To investigate the subcellular locations of maize COI proteins, on gene of each closely related pair (*COI1a*, *COI1c*, and *COI2a*) was fused to *GFP* and transiently expressed in *Nicotiana benthamiana*. Two hours after spraying the plants with MeJA, the majority of COI2a protein was fully localized in the nucleus, similar to what is observed with COI in Arabidopsis (Withers et al., 2012). By contrast, COI1a and COI1c showed nucleo-cytosolic partitioning and were partly in droplet-like subcellular structures in the cytosol (Figure 2). Nucleo-cytosolic partitioning of transcription factors and their regulators, an emerging field in developmental biology, has essential functions in hormonal regulation of plant physiology (Jing et al., 2022; Powers et al., 2019). To investigate whether a different ligand can increase the nuclear localization of maize COI1A and COI1B, we tested the effect of the oxylipin 12-oxo-phytodienoic acid (OPDA), the immediate precursor of JA, which is an alternative ligand for COI (Jimenez Aleman et al., 2022; Varsani et al., 2019). However, OPDA did not increase the extent of COI1 nuclear localization under the tested conditions (Figure S3).

**Figure 2.**
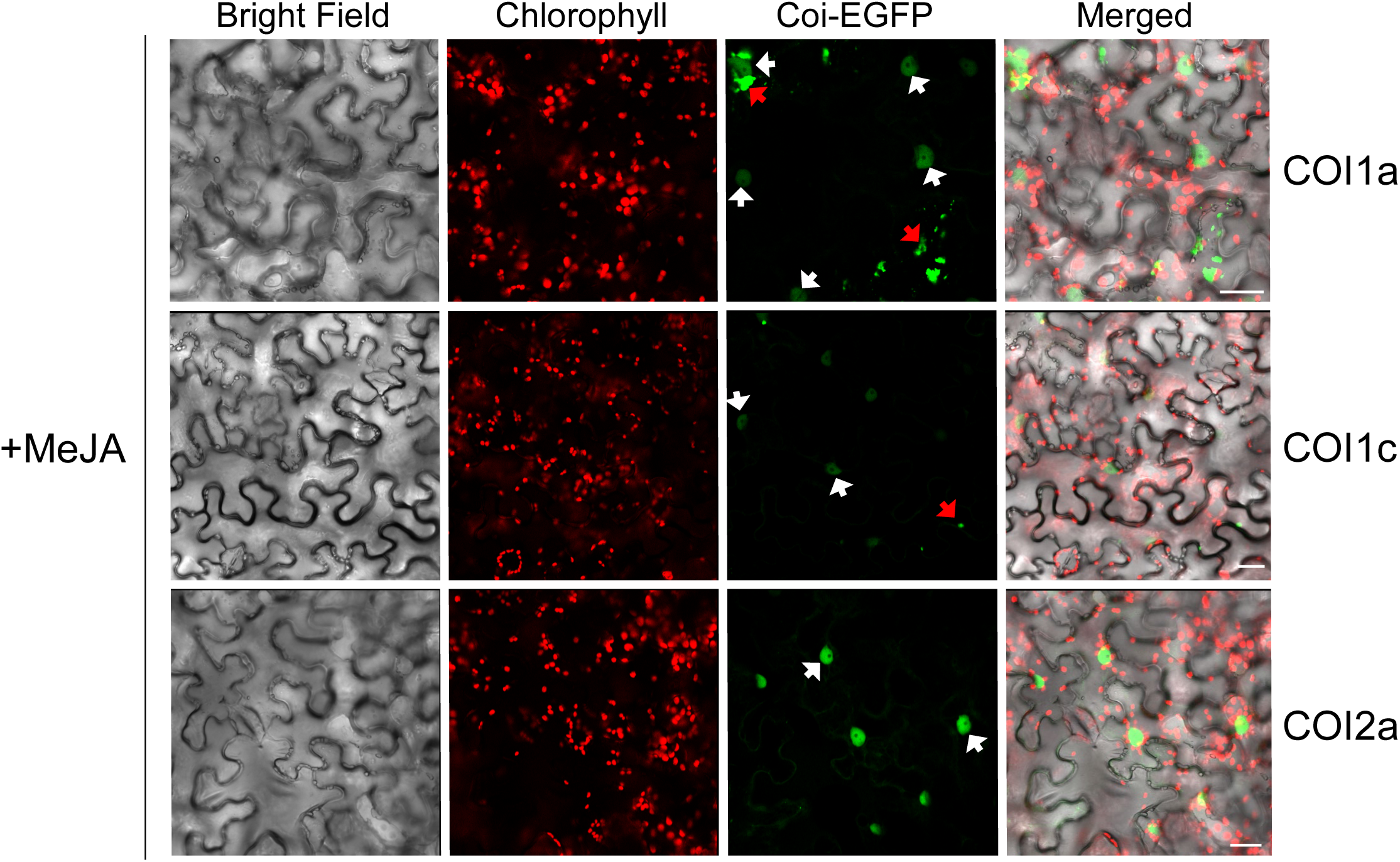
Subcellular locations of the maize COI1a, COI1c, and COI2a proteins fused to enhanced green fluorescent protein (EGFP) show that COI2a is more targeted to the nuclei than the two COI1 proteins. Confocal images were taken at 48 h after infiltrating plasmids expressing the COI-EGFP fusion proteins into the *Nicotiana benthamiana*. Leaves were sprayed with MeJA two hours before imaging. White arrows indicate the nuclei, and red arrows represent cytosolic condensates. Scale bars = 25 μm.

The canonical functions of COI proteins include binding to JAZ repressors. Using a bimolecular fluorescence complementation (BiFC) vector system, we co-expressed *COI1a*, *COI1c*, and *COI2a*, fused to the C-terminal half of the *YFP*, together with fifteen members of the maize *JAZ* gene family fused to the N-terminal half of the *YFP*. This showed that, whereas COI2a strongly interacted with almost half of the maize JAZ proteins in *N. benthamiana* leaf nuclei (Figure 3, and S4), neither COI1a nor COI1c showed a reproducible interaction with any JAZ proteins after induction by treatment with MeJA (Figures 3, and S4). An independent repeat of this experiment showed similar results (Figure S5). We tested both OPDA and 9-hydroperoxy-10,12,15-octadecatrienoic acid (9-HPOT), a metabolic precursor of the death acids, which are positional isomers of the JA-like oxylipins (Vellosillo et al., 2013; Christensen et al., 2015), and found that they did not change COI-JAZ interactions (data not shown). Together, these experiments indicate that, unlike COI2, COI1A and COI1B do not efficiently bind to JAZ in the presence of MeJA or potentially JA conjugates that were derived from MeJA *in planta*, suggesting that the maize COI1 proteins fully or partially lost their canonical functions after their divergence from the COI2 proteins.

**Figure 3.**
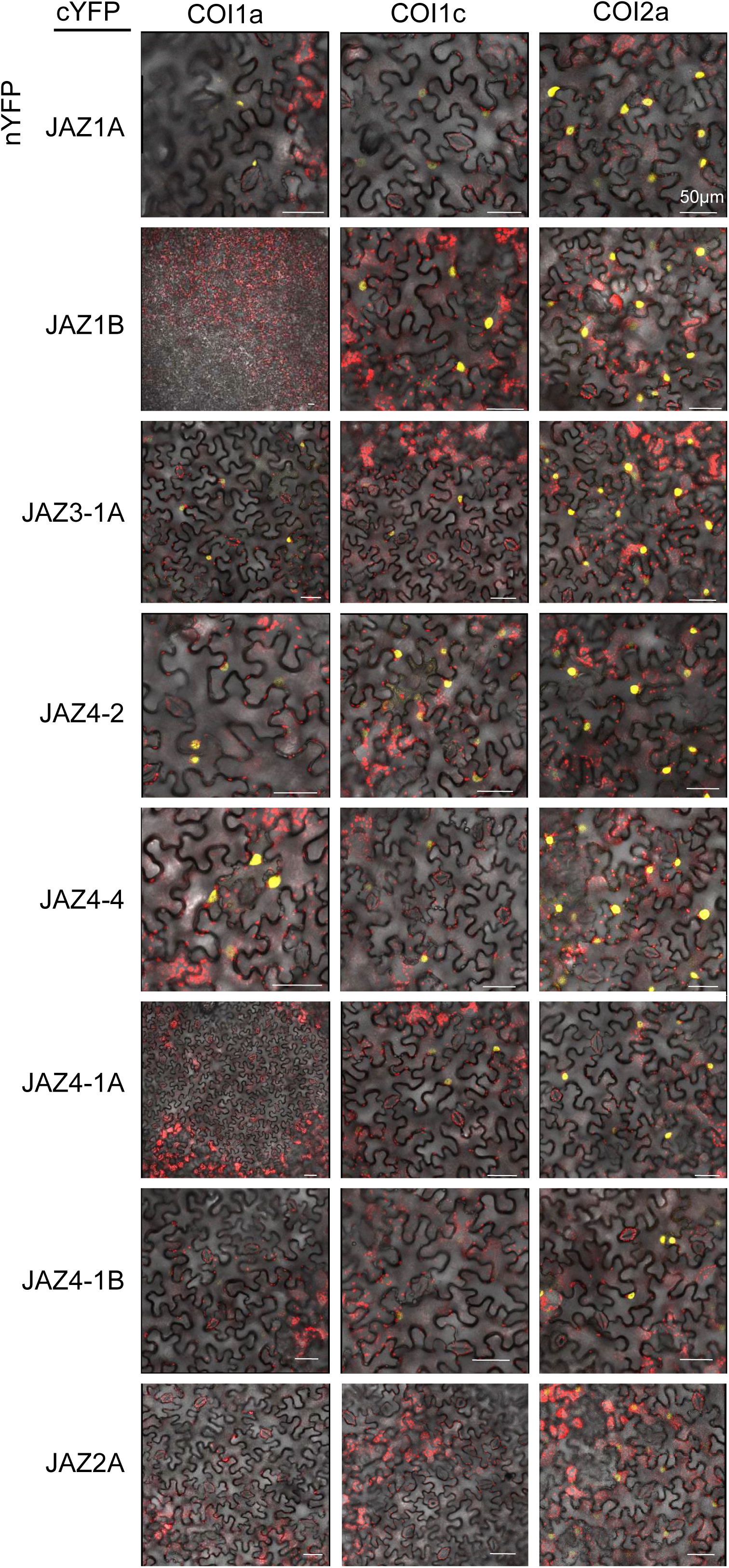
Bimolecular fluorescence complementation (BIFC) between the maize COI1a, COI1c, or COI2a and eight JAZ proteins shows that JAZ proteins have a higher affinity for COI2 than COI1. Scale bars = 50 μm. These experiments were repeated independently with similar results (Figure S5).

### Generation of higher-order mutants of the maize *COI* genes

To investigate the *in vivo* functions of the maize COI proteins, we identified *Ds* and *Mu* transposon insertions in the genetic background of maize inbred line W22 (Springer et al., 2018). There was a pre-existing *Ds* transposon insertion mutation in the *COI1d* gene (Vollbrecht et al., 2010; Ahern et al., 2009), and we remobilized other *Ds* transposon insertions to create mutations in *COI1a*, *COI1c*, and *COI1b*. We obtained *Mu* transposon-insertion mutations of the *COI2* genes from the UniformMu Transposon Resource (Liu et al., 2016a; Settles et al., 2007). Genotyping of the mutants by PCR identified the position of each transposon insertion (Figure 4A; Table S3). All of these transposon insertions are in the coding regions of their respective genes, most likely causing loss-of-function mutations due to in-frame stop codons in the transposon sequences or aberrant splicing (Simon and Starlinger, 1987).

**Figure 4.**
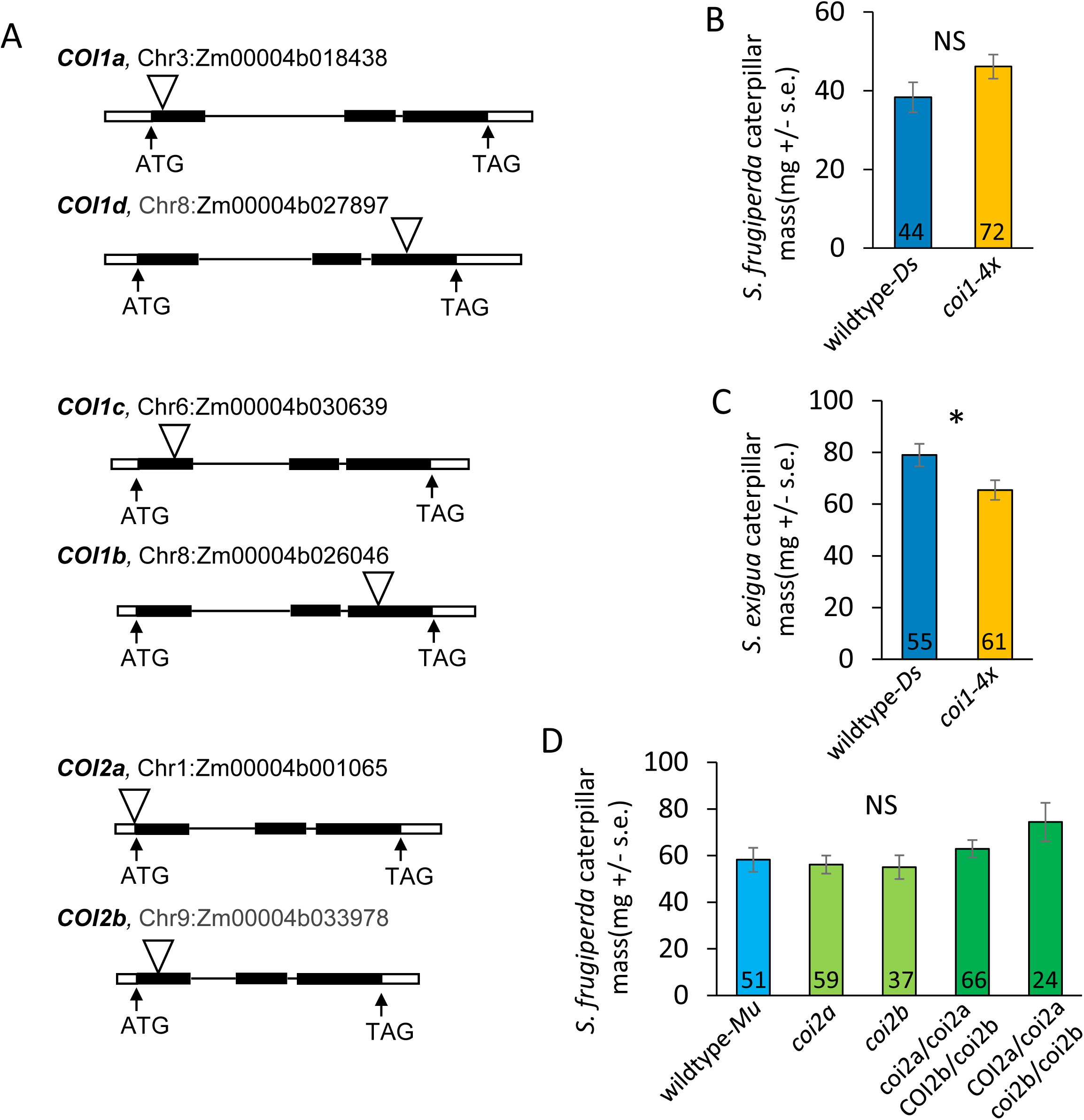
Caterpillar growth on *coi* mutant and wildtype maize. (A) Locations of *Ds* transposon insertions in *COI1* genes and *Mu* transposon insertions in *COI2* genes are marked with triangles. Insertions are located at 140, 2682, 344, 2342, –8, and 198 bp from the start codons of *COI1a*, *COI1d*, *COI1c*, *COI1b*, *COI2a*, and *COI2b*, respectively. Black bars represent exons, thin lines are introns, and white bars are non-coding regions of the genes. (B) Mass of ten-day-old *Spodoptera frugiperda* (fall armyworm) caterpillars on wildtype and *coi1-4x* maize. No significant difference (P > 0.05, *t-* test). (C) Mass of ten-day-old *Spodoptera exigua* (beet armyworm) caterpillars on wildtype and *coi1-4x.* *P < 0.05, *t-*test. (D) Mass of ten-day-old *S. frugiperda* on wildtype and *coi2* mutants. No significant difference (NS, P > 0.05), Dunnett’s test relative to wildtype-*Mu*. All data are mean +/-s.e., numbers in bars indicate sample sizes.

Homozygous *coi* mutants were intercrossed to generate a set of higher-order mutants. This included a homozygous *coi1a coi1d* double mutant, a homozygous *coi1b coi1c* double mutant, and a homozygous quadruple mutant of all four *COI1* genes, hereafter called *coi1-4x*. The *COI2* combinations included a *coi2a coi2b* double mutant that was homozygous for *coi2a* and heterozygous for the *coi2b* (*coi2a/coi2a COI2b/coi2b*) and a double mutant that was heterozygous for *coi2a* and homozygous for *coi2b* (*COI2a/coi2a coi2b/coi2b*).

### Knockout of both *COI2* genes leads to pollen lethality

As has been reported previously for CRISPR/Cas9 mutants of these genes (Qi et al., 2022), a *coi2a coi2b* double knockout is pollen lethal. We were not able to obtain a homozygous double mutant by self-pollination of *coi2a/coi2a COI2b/coi2b* or *COI2a/coi2a coi2b/coi2b.* Genotyping of the progeny from these self-pollinations showed a 1:1 ratio of the parental genotype and homozygous single mutants. This ratio is expected if a double knockout *COI2* genes results in gametophyte lethality rather than embryo lethality (Figure S6). To investigate whether *coi2a coi2b* is lethal to pollen or egg cells, we performed test crosses between the *coi2a/coi2a COI2b/coi2b* and *COI2a/coi2a coi2b/coi2b* and wildtype W22, with the wildtype being either the pollen or the egg donor. Genotyping of the progeny indicated that, whereas *coi2a*/*coi2b* eggs are viable and can be transmitted to the next generation, the *coi2a coi2b* pollen genotype is lethal (Figure S7). The tissue-specific abundance of the *COI2a* transcripts in anthers and the *COI2b* gene in mature pollen (Figure S8, plotted using the Maize Genome Database, www.maizegdb.org) and the COI2a protein in pollen (Figure S9; data from (Walley et al., 2016)) further suggested that the COI2 proteins contribute to pollen development.

### Resistance to lepidopteran herbivory is not compromised in *coi* mutants

Perception of JA-Ile by COI proteins is associated with the induction of herbivore defenses in other plant species. However, *Spodoptera frugiperda* (fall armyworm) did not grow larger on *coi1-4x* (Figure 4B), and *Spodoptera exigua* (beet armyworm) grew less well on *coi-4x* than on wildtype W22 plants (Figure 4C). This suggested that the COI1 proteins by themselves do not play a major role in maize defense against insect herbivores. On *coi2a/coi2a COI2b/coi2b* and *COI2a/coi2a coi2b/coi2b* mutant lines, *S. frugiperda* growth was not significantly improved relative to wildtype parental lines (Figure 4D). However, as it was not possible to obtain homozygous *coi2a coi2b* double mutants, and a single functional *COI2* gene may allow defense induction, residual COI activity may be sufficient to limit herbivory.

### *COI1* quadruple mutants have shortened internodes

When growing the *coi1-4x* mutant line, we noticed that the plants are shorter than wildtype W22 and the *coi1* double mutants (*coi1a coi1d* and *coi1b coi1c*) derived from the same crosses. At ten days post-germination, the *coi1-4x* mutants are similar in size to wildtype (Figure S10A). However, whereas the first two leaves were longer in *coi1-4x* mutants than in wildtype, the third leaves were significantly shorter, marking the onset of a developmental delay in the mutant (Figure 5A,B). The *coi1-4x* mutants were almost half of the size of wildtype W22 and both of the *coi1-2x* mutants at 20 days (Figure S10B, C) and 60 days (Figure 5C,D) post-germination. The reduced height at 60 days post-germination was primarily due to shortened internodes in the *coi1-4x* plants relative to the corresponding internode lengths in *coi1a coi1d* and *coi1b coi1c*, and wildtype lines derived from the same crosses (Figure 5E,F).

**Figure 5.**
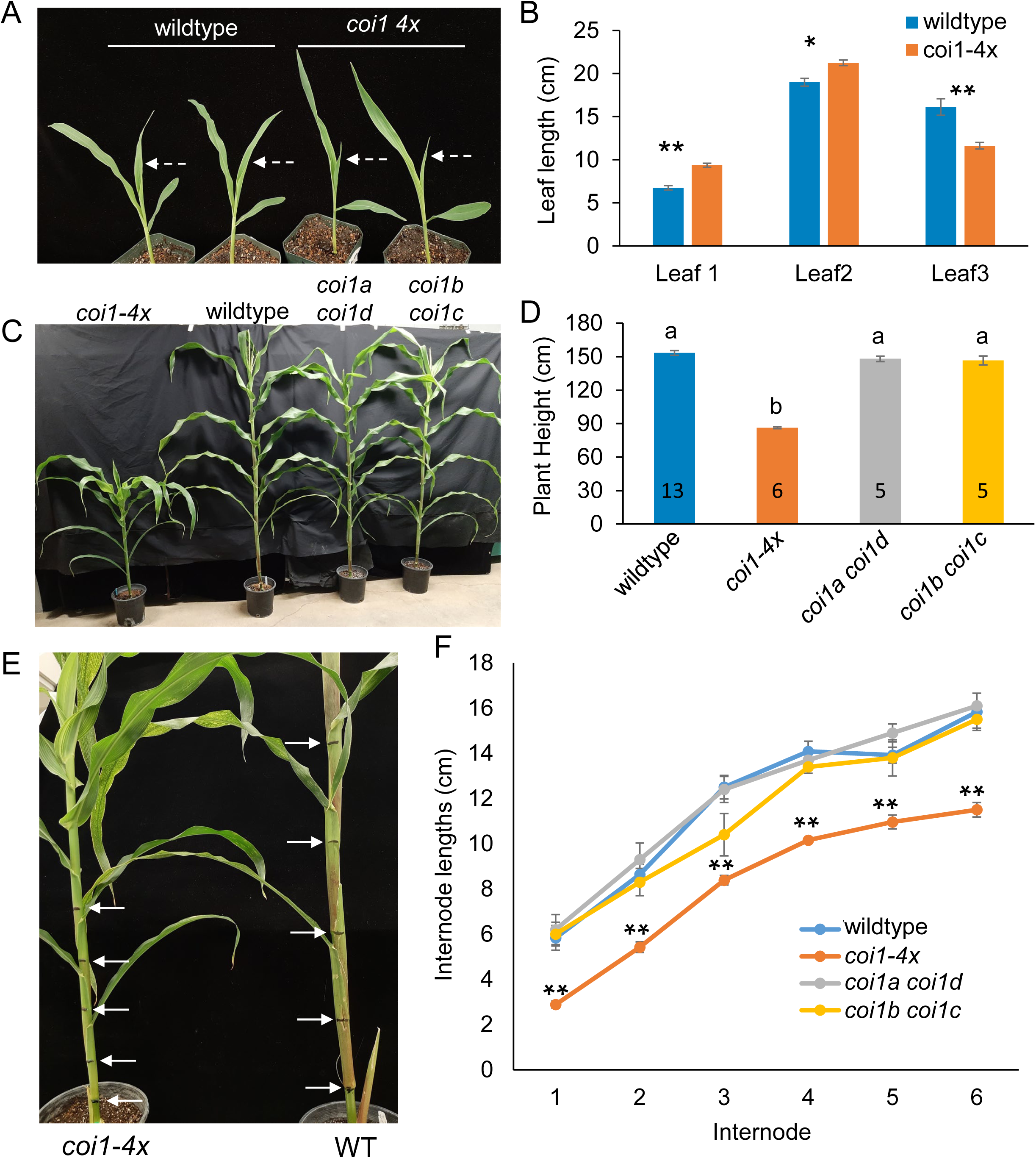
*coi1-4x* has impaired growth relative to the corresponding double mutants, *coi1a coi1d* and *coi1b coi1c*, and wildtype inbred line W22 maize. (A,B) Leaf lengths of *coi-4x* and wildtype (WT) ten days after germination, N = 4, mean ± se, two-tailed Student’s *t*-test, *P < 0.05, **P < 0.01. (C,D) Plant heights at 60 days after germination. Numbers in bars = sample sizes, mean +/-s.e., letters indicate significant differences, P < 0.05, ANOVA followed by Tukey’s HSD test. (E,F) Internode lengths were compared between four genotypes at 60 days post-germination. *coi1-4x* N = 13, wildtype N = 6, *coi1a coi1d* and *coi1b coi1c* N = 5, mean +/-s.e., **P < 0.01, Dunnett’s test relative to wildtype.

### The *coi1-4x* mutant has reduced leaf micronutrients and photosynthetic activity

The consistently shorter growth of the *coi1-4x* mutant (Figure 5) was accompanied by a striped-leaf phenotype that persisted throughout the life of the plant (Figure 6A). As striped leaves can indicate a mineral nutrient deficiency (Mattiello et al., 2015; Curie, Catherine et al., 2001; Foy and Barber, 1958; Thoiron et al., 1997), we measured macro-and microelements in the leaves of *coi1-4x*, *coi1a coi1d*, *coi1b coi1c*, and wildtype W22 using inductively coupled plasma atomic absorption emission spectroscopy (ICP-MS). This showed a decrease in leaf iron, manganese, copper, and zinc in the *coi1-4x* mutant relative to the double mutants and wildtype 20-day-old plants (Figure 6B). Similar reductions in iron, manganese, copper, and boron in *coi1-4x* were seen in 13-day-old plants in a separate experiment (Figure S11A). The levels of leaf macro elements showed only a few differences between the *coi1-4x* mutants and the other lines, a decrease in phosphorous at 20 days (Figure 6C) and an increase in potassium at 13 days (Figure S11B) after germination.

**Figure 6.**
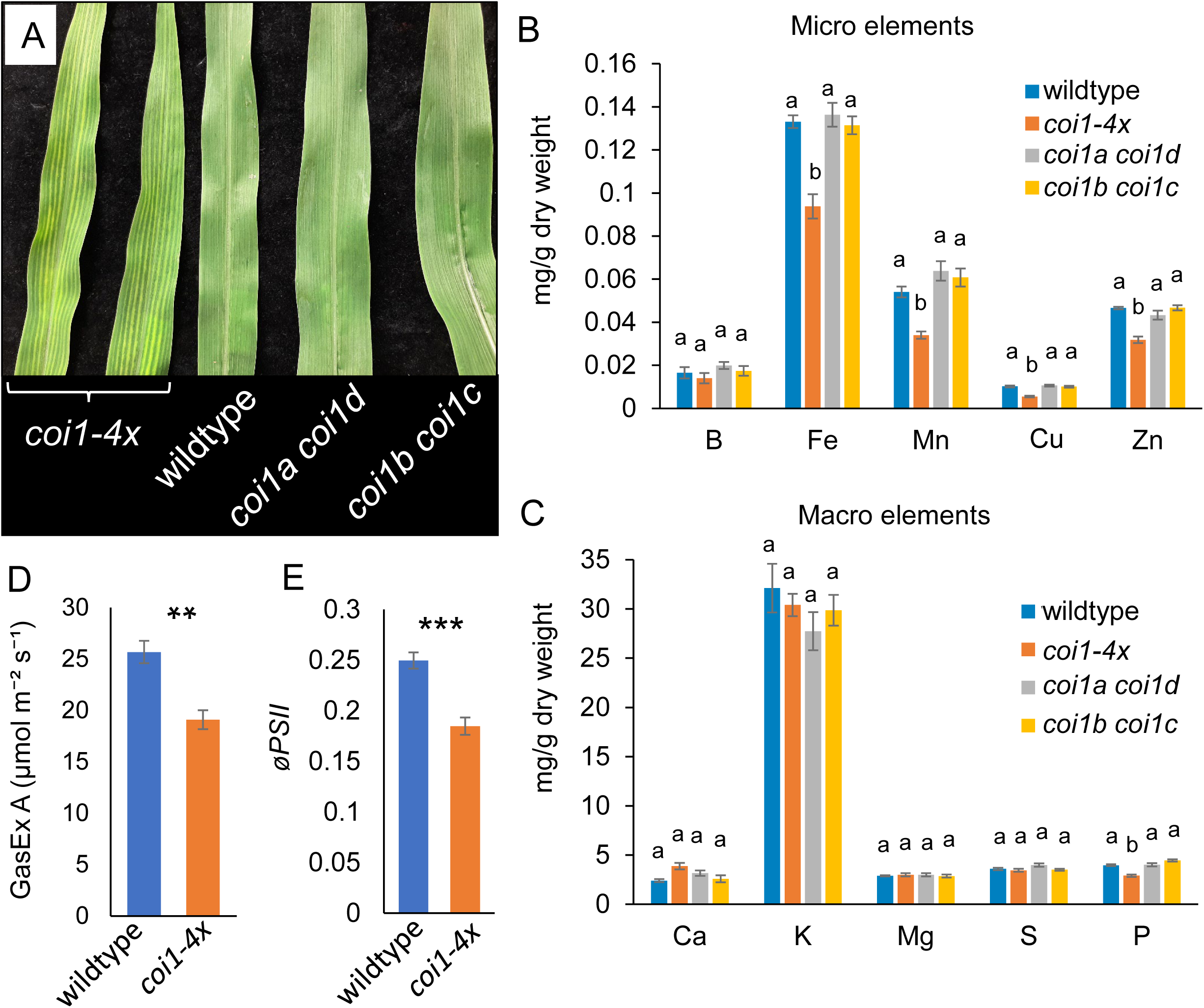
*coi1-4x* plants have striped leaves, decreased microelement levels, and reduced photosynthesis. (A) The striped leaf phenotype of the *coi1-4x* at 20 days post-germination compared to corresponding leaves from the double mutants and wildtype W22. (B) Microelements and (C) macroelements in 20-day-old seedlings, N = 7, mean +/-s.e., letters indicate differences (P < 0.05) using Tukey’s HSD test. (D) Leaf CO_2_ assimilation rate (GasEX A) at 400 µmol mol⁻¹ CO_2_ and (E) the quantum yield of the photosystem II phytochemistry (*øPSII*) at 2000 μmol m^-2^ s ^-1^ of actinic light (B) were measured at 20 days after germination, respectively. Mean ± s.e., n = 20, two-tailed Student’s *t*-test, \*\**P* < 0.01, \*\*\**P* < 0.001.

Because the striped leaf phenotype can also indicate photosynthetic deficiency, we compared the carbon assimilation efficiency and photosynthesis quantum yields between *coi1-4x* and wildtype. This showed a decrease in both carbon assimilation and quantum yield in the *coi1-4x* mutant at 20 days after germination (Figure 6D, E). In a separate experiment, a similar photosynthetic defect was observed in 28-day-old plants (Figure S11C, D).

### Differential gene expression in *coi* mutants, with or without MeJA elicitation

We used transcript profiling to identify differentially expressed genes in the *coi* mutants compared to wildtype plants, with and without MeJA elicitation. The 13 tested genotypes consisted of six *coi* single mutants, *coi1a coi1d*, *coi1b coi1c*, *coi1-4x*, double homozygote-heterozygote combinations of the *coi2* mutations, and two corresponding inbred line W22 wildtype lines (for the *Ds* and *Mu* insertion line crosses, respectively). Leaf tissue for gene expression assays was taken from 5-9 biological replicates of the thirteen genotypes, each sprayed with MeJA or buffer as a control. Reads per million mapped reads (RPM) were calculated for each gene, and *P* values for each between-genotype or between-treatment pairwise comparison were calculated for 40,690 annotated genes in the maize W22 genome (Springer et al., 2018). In one or more of 73 pairwise comparisons that were quantified among the 26 tested conditions (13 genotypes +/-MeJA; Supplemental Dataset S1), there were 13,365 genes with differential expression (adjusted *P* <0.05; Supplemental Dataset S2). A heatmap based on this list of differentially expressed genes showed that most expression differences were due to induction or repression by MeJA (Figure 7).

**Figure 7.**
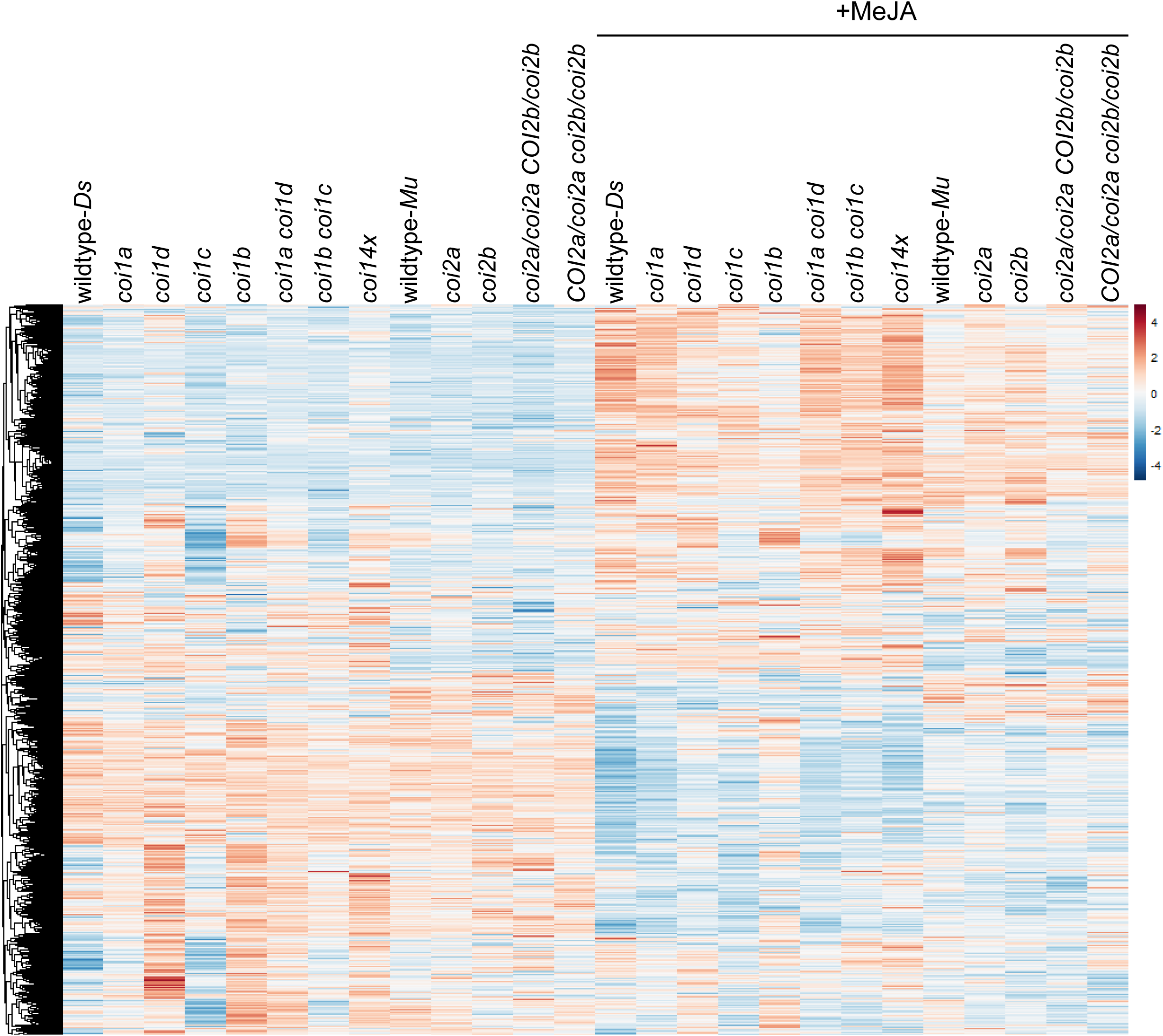
Heatmap of gene expression for 13,365 genes that were differentially regulated between two or more genotypes or between mock and methyl jasmonate (MeJA) induction. Color ranges from blue (minimum) to red (maximum) reads per million bp for each gene.

### Transcripts accumulating to lower levels in *coi1-4x* encode components of photosynthetic machinery and cell wall metabolism

As we were interested in the short stature and photosynthetic defects that differentiate the *coi1-4x* mutant from wildtype W22 and other *coi* mutant lines (Figures 5 and 6), we searched specifically for genes that were differentially expressed in *coi1-4x* relative to *coi1a coi1d*, *coi1b coi1c*, *coi2a/coi2a COI2b/coi2b*, *COI2a/coi2a coi2b/coi2b*, and wildtype W22, with or without MeJA treatment. This identified 25 genes that were expressed at a significantly downregulated (Figure 8A and Table S4) and 68 genes that were significantly upregulated (Figure 9A and Table S5) in *coi1-4x* relative to the other genotypes.

**Figure 8.**
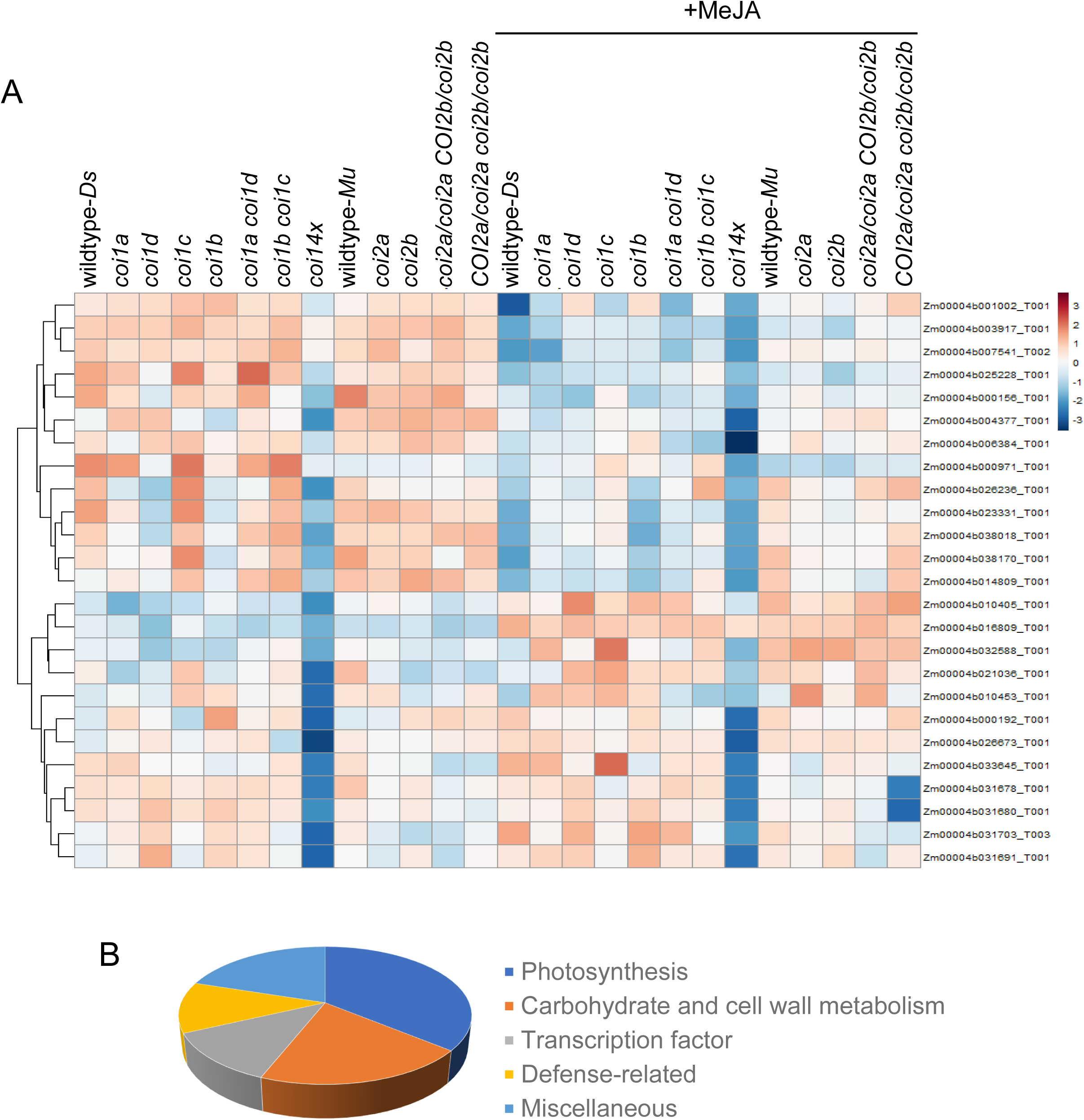
Genes down-regulated in the *coi1-4X* mutant encode two main groups of proteins, involved in C_4_ photosynthesis and carbohydrate and cell wall metabolism. (A) Heatmap showing downregulated genes in *coi1-4x* relative to other genotypes, with or without methyl jasmonate (MeJA) treatment. The reads are transformed by log(1+expression) prior to clustering. (B) Down-regulated genes in *coi1-4x* were categorized into five functional groups. The list of the genes, color-coded by their functional group, as well as ordered based on their position on the heatmap, is presented in Table S4.

**Figure 9.**
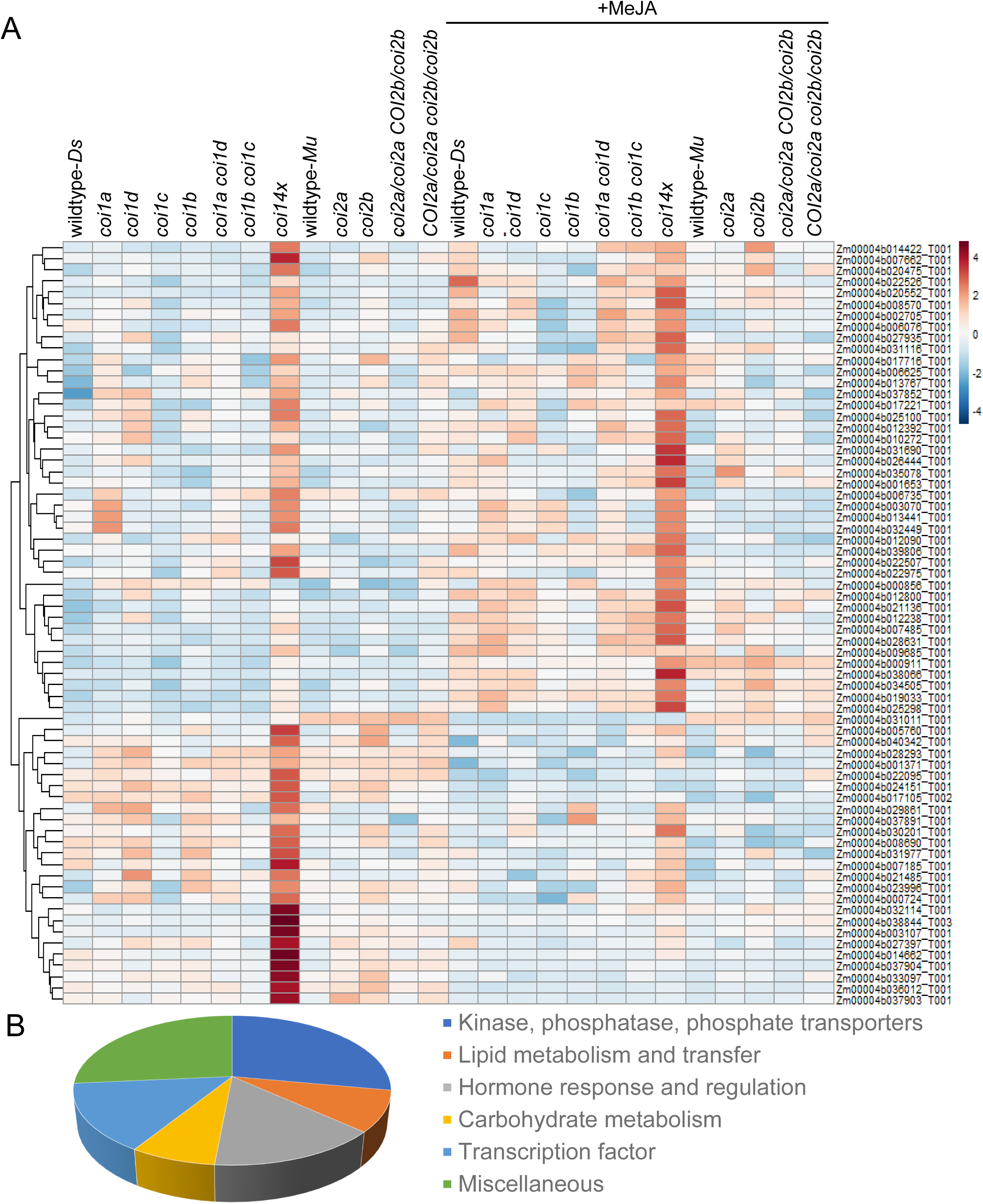
Genes that are up-regulated in the *coi1-4x* mutant encode proteins involved in phosphate regulation, lipid and carbohydrate metabolism, hormone regulation, and transcription. (A) Heatmap showing upregulated genes in *coi1-4x* relative to other genotypes, with or without methyl jasmonate (MeJA) treatment. The reads are transformed by log(1+expression) prior to clustering. (B) Up-regulated genes in *coi1-4x* were categorized into six functional groups. The list of the genes, color-coded by their functional group, as well as ordered based on their position on the heatmap, is presented in Table S5.

The downregulated genes in the *coi1-4x* mutant are clustered in two groups in the heatmap dendrogram (Figure 8A), based on whether they are upregulated (top half) or downregulated (bottom half) by MeJA in the other twelve maize lines. Genes that were repressed by MeJA treatment were expressed at a low level in *coi1-4x*, even prior to MeJA elicitation. Conversely, genes that were induced by MeJA in other lines are still expressed at a relatively low level in *coi1-4x* after MeJA.

Among the downregulated genes in the *coi1-4x* mutant, 72% encode bundle sheath or mesophyll-abundant transcripts (Table S6), based on previously published cell-type-specific transcriptome data (Li et al., 2010). We grouped the downregulated gene list (Figure 8B and Supplemental Dataset S1) based on functional annotations in the Maize Genome Database (www.maizegdb.org) and homologs in The Arabidopsis Information Resource (TAIR, www.arabidopsis.org). Genes involved in photosynthesis, carbon fixation, and cyclic electron transport constituted almost one-third of the downregulated genes in the *coi1-4x* mutant, consistent with this mutant’s decrease in carbon assimilation and photosystem II quantum yield (Figure 6D, E). These included genes encoding Calvin cycle enzymes (Rubisco small subunit and phosphoribulokinase1 (PRK1)), C_4_ photosynthesis enzymes (phosphoenolpyruvate carboxykinase1 (PEPCK1), alpha carbonic anhydrase, orthophosphate dikinase2 (PPDK2), and NAD(P)H-quinone oxidoreductase subunit U (NadhU)), fibrillin 4, which is associated with the photosystem II light-harvesting complex, thylakoids, and plastoglobules (Singh et al., 2010), and a chloroplast co-chaperonin. NadhU is an essential part of the NAD(P)H dehydrogenase complex (NDH), which performs cyclic electron transport in chloroplasts by transferring electrons from ferredoxin to plastoquinone (Yamamoto et al., 2011), and fibrilins have essential roles in the biosynthesis of plastoquinone, light acclimation, and sulfur metabolism (Kim et al., 2015; Lee et al., 2020).

Another group of downregulated genes encoded proteins involved in cell wall metabolism, carbohydrate breakdown, and reactive oxygen species (ROS) scavenging. This group included those encoding peroxidase 52 and a cinnamyl alcohol dehydrogenase (CAD), which are both involved in lignin biosynthesis, a copper/zinc superoxide dismutase 1 (CSD1), Sugary 1 (SU1, a pectin methylesterase inhibitor), which has a role in amylopectin biosynthesis, and an invertase involved in sucrose breakdown.

A small group of transcription factors, including the HAP5-transcription factor (*NF-YC4*), which is involved in GA and abscisic acid (ABA)-activated signaling pathways (37), *MYC70*, a bHLH-transcription factor, and a cell growth defect protein with unknown function, were among the genes that were downregulated in the *coi1-4x* mutant. We did not find any downregulated genes with a known role in homeostasis, regulation, or transport of the micronutrients that could explain the specific deficiency of these elements in the *coi1-4x* mutant (Figure 6B). However, it is possible that the downregulation of some of the cell wall and lignin-related proteins, such as *CAD*, which have been implicated in zinc and iron homeostasis, might be the underlying cause of the microelement deficiency in the *coi1-4x* mutant.

Only a few defense-associated genes were downregulated in the *coi1-4x* mutant (Figure 8), including those encoding terpene synthase 19 (TPS19), halotolerant determinant 3-like protein (HAL3A, a heme-binding protein with a putative role in terpenoid metabolism), a ras-group-related LRR protein, an isoflavone reductase, and a serine-type endopeptidase. Expression of well-studied JA-induced maize defense genes, including benzoxazinoid biosynthesis genes (Figure S12 and Table S7), terpene synthase genes (other than *TPS19*) (Figure S13 and Table S8), and *JAZ* genes (Figure S14 and Table S9), was not altered in the *coi1-4x* mutant relative to the other genotypes. This is consistent with the observation that resistance to lepidopteran herbivory was not compromised in the *coi1-4x* mutant compared to W22.

### Genes with higher transcript abundance in *coi1-4x* are mainly involved in phosphate metabolism, cell wall modification, and transcription regulation

A dendrogram and heatmap of genes that are expressed at a higher level in the *coi1-4x* mutant than in the other genotypes (Figure 9A and Table S5) also had two clusters, genes that are induced (top part) or repressed (bottom part) by MeJA. These genes were expressed at a higher level in *coi1-*4x, with or without MeJA treatment. Altogether, 29% of the genes encode bundle sheath or mesophyll-abundant transcripts (Table S10) (Li et al., 2010). When these upregulated genes were grouped by their functional annotations, the most abundant group comprised those encoding kinases, phosphatases, and a few transporters involved in the transport of phosphate and other molecules, some of which are implicated in phosphate homeostasis (Figure 9B and Table S5). A smaller group of upregulated genes had roles in lipid and phospholipid or glycolipid metabolism. Noticeably in this group, glycerophosphodiester phosphodiesterase 2 (*GPX2*) and monogalactosyldiacylglycerol synthase type c (*MGD3*) catalyze lipid metabolism during phosphate starvation. Another relatively large group of upregulated transcripts included genes that were shown to be hormone-regulated in prior research. Most of these genes, including two PP2C phosphatase genes, were induced by abscisic acid. This group also included those encoding Friendly Mitochondria (FMT), non-responding to oxylipins 38 (NOXY38), and lipoxygenase 12, a 9-LOX that is the main enzyme involved in the biosynthesis of the JA stereoisomers called death acids (Christensen et al., 2015). FMT regulates mitochondrial distribution, fusion, and quality control in Arabidopsis, but has also been shown to perceive 9-LOX-derived oxylipins (Vellosillo et al., 2013, 2007).

### COI1 proteins function in GA signaling

The two main downregulated gene groups (photosynthesis and cell wall metabolism) in the *coi1-4x* mutant are induced by the GA signaling pathway (Ranwala and Miller, 2008; Chen et al., 2020; Falcioni et al., 2018). Moreover, the short stature and shortened internodes of the *coi1-4x* mutant are similar to what is observed in constitutively active, dominant *DELLA* mutants, which also have a dwarf phenotype (Lawit et al., 2010; Ueguchi-Tanaka et al., 2008; Cassani et al., 2009; Fujioka et al., 1988; Gale and Marshall, 1973; Strader et al., 2004). Therefore, we hypothesized that, unlike Arabidopsis and rice *coi* mutants, which are elongated relative to the respective wildtype plants (Xu et al., 2002; Machado et al., 2017; Yang et al., 2012; Inagaki et al., 2022), the maize *coi1-4x* mutant is hyposensitive to GA signaling. To test this hypothesis, we used an antibody against rice DELLA (SLR1) (Ueguchi-Tanaka et al., 2008) to detect the maize DELLA protein and compare its abundance in the leaves of 20-day-old *coi1-4x* mutant and wildtype W22 maize seedlings. A protein band migrated at the predicted size of DELLA, over-accumulated in the mutant plants, suggesting that DELLA is more stable in the *coi1-4x* mutant (Figure 10). To confirm that this protein band is DELLA, we treated the 5-day-old *coi1-4x* and wildtype seedlings germinated in water-filled pouches, with 0.02% GA and 0.02% MeJA for two days. Immunoblot analysis of the leaves from these seedlings showed that DELLA was reduced and increased after GA and MeJA treatments, respectively (Figure S15). The level of DELLA, however, was not different between the *coi1-4x* and wildtype seedlings in this experiment. This was expected because the wildtype and *coi1-4x* seedlings at this early age and after germination in water, show height and leaf phenotypes that are similar to wildtype plants.

**Figure 10.**
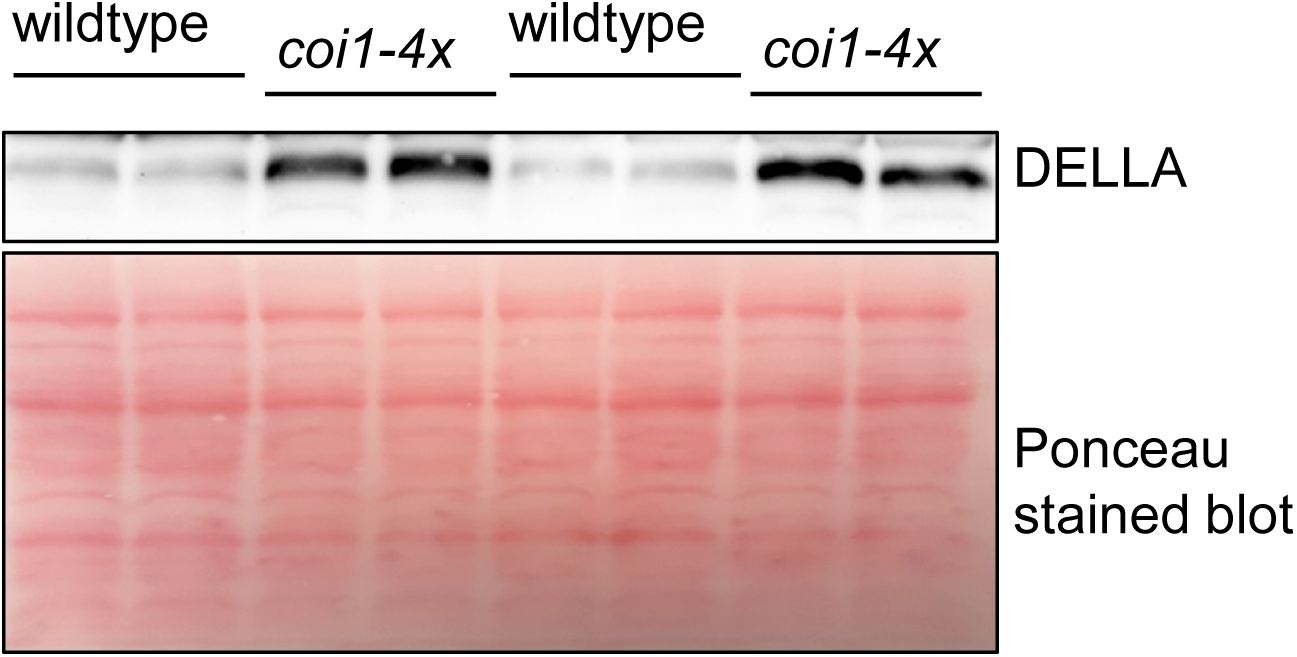
Immunoblot analyses of maize DELLA proteins. The leaf total proteins from equal surface area (similar weight) of *coi1-4x* and wildtype inbred line W22 leaves were analyzed by probing with the rice SLR1 antibody. The membrane was stained with Ponceau S as the loading control.

To further investigate the effect of COI1 on DELLA, we examined protein-protein interactions. We did not observe any interaction between COI proteins and DELLA using BIFC. However, co-expression of *DELLA-RFP* with *COI-GFP* in *N. benthamiana* lent support to the hypothesis that these proteins may interact directly or indirectly through a larger protein complex. While DELLA-RFP was highly expressed and fully localized in the nuclei when expressed alone or with GFP, it disappeared upon co-expression with any of the three tested COI proteins, COI1a, COI1c, or COI2a (Figure 11). Overall, less than 1% of the nuclei contained DELLA-RFP in the presence of COI. However, we could still detect DELLA-RFP outside of the nuclei, colocalized with the COI1-GFP in the unknown cytosolic structures (Figure S16). We previously observed these structures with COI1-GFP alone (red arrows in Figure 2).

**Figure 11.**
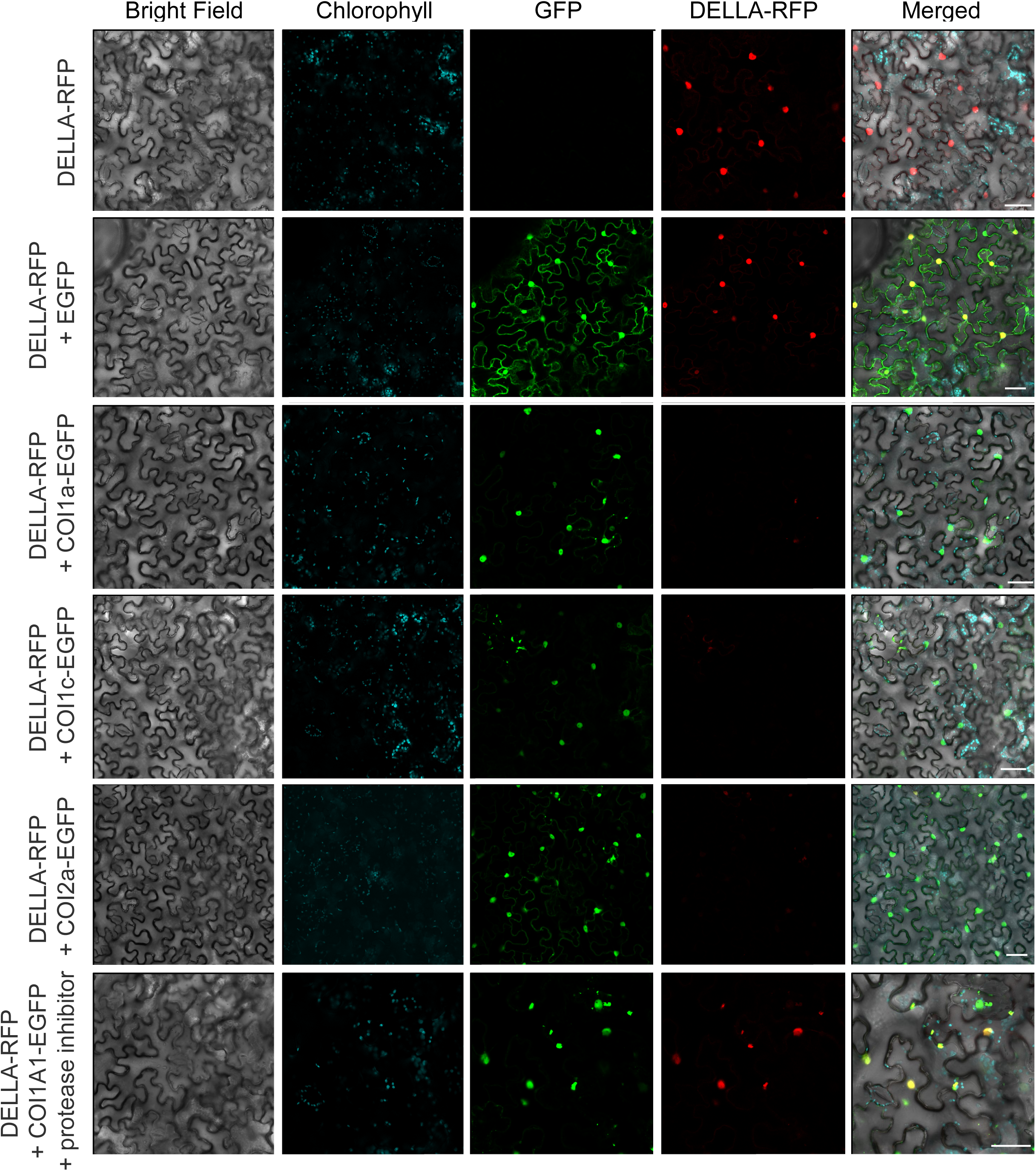
Maize DELLA (DWARF9) disappears from the nuclei upon coexpression with the maize COI proteins. Confocal images were taken after expressing genes in *Nicotiana benthamiana* leaves. Labels on left indicate infiltrated genes for each row of images. Protease inhibitor (bortezomib) was co-infiltrated in the bottom row. Scale bars = 50 μm.

COI is the F-box domain protein component of an E3-ligase complex that polyubiquitinylates JAZ repressors and sends them for degradation by the 26S proteasome. We hypothesized that the maize COI proteins also participate directly or indirectly in the assembly of E3-ligase complexes that degrade DELLA. Therefore, we infiltrated leaves co-expressing DELLA-RFP and COI1a-GFP with the proteasome inhibitor bortezomib (BTZ) to test this hypothesis. BTZ treatment restored DELLA in 60% of the nuclei (Figure 12A). Immunoblotting of the total proteins extracted from the leaves with SLR1 antibody, showed that BTZ treatment of the leaves that co-expressed DELLA-RFP with COI1a, restored the level of DELLA-RFP from 25% to more than 50% of the controls (expressing DELLA-RFP alone or with GFP) (Figure 12B). The similar BTZ-mediated restoration for the COI1c was from more than 50% to 100%. Together, these experiments confirmed that COI directly or indirectly targets DELLA for degradation by proteasomes.

**Figure 12.**
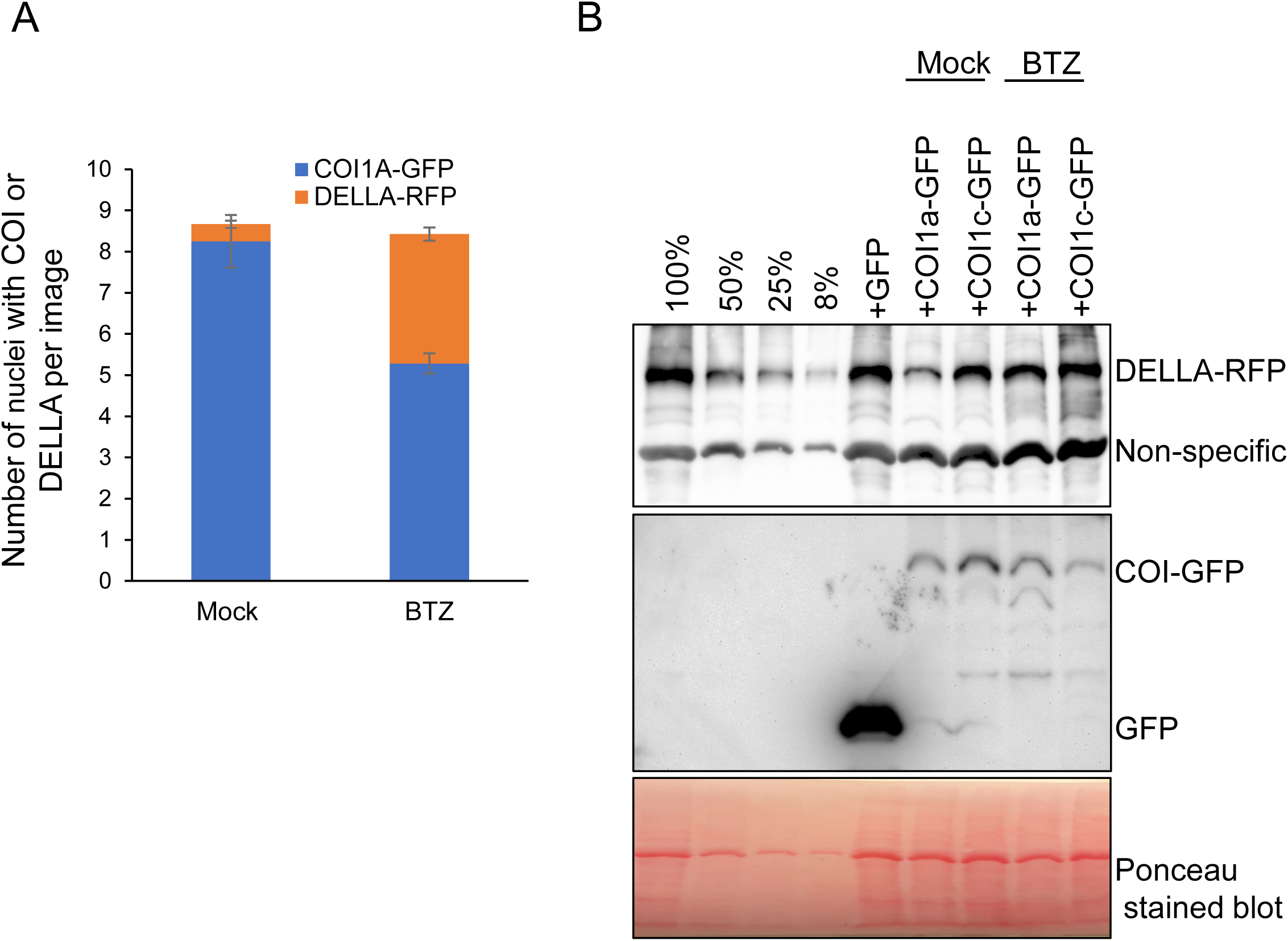
Expression of the maize COI1 proteins leads to proteasome-dependent degradation of maize DELLA in *N. benthamiana*. (A) Nuclei showing the presence of the COI1a-EGFP and or DELLA-RFP were counted on confocal images from an experiment similar to Figure 11, with and without protease inhibitor. (B) Leaf tissue from the *N. benthamiana* plants expressing DELLA-RFP alone or with EGFP, COI1a-EGFP, or COI1c-EGFP (similar to those used in Figure 11) were used for immunoblot analyses. Membranes were probed with the rice SLR1 and GFP antibodies. The first membrane was stained with Ponceau S as the loading control. Non-specific binding of the SLR1 antibody to an unrelated protein was used as an extra loading control.

### Effects of Exogenous Hormones on the *coi1-4x* Growth Phenotypes

To further investigate the effects of JA and GA signaling on the *coi1-4x* and wildtype maize, one-week-old seedlings of both genotypes were watered with either 0.02% MeJA or 0.02% GA for three weeks. Three weeks after MeJA treatment, *coi1-4x* leaves were chlorotic, with a yellow color (Figure 13A). By contrast, wildtype leaves only showed some stripes during the first week of treatment but then became green again. After two weeks, the height of MeJA-treated wildtype plants was reduced by 53% relative to mock-treated controls (Figure 13B, D). Growth reduction of the *coi1-4x* mutant was less severe (35%), suggesting that the mutant line was less sensitive to the growth-inhibitory effect of the MeJA. We attributed this continued MeJA sensitivity to the presence of the two functional *COI2* genes in the *coi1-4x* mutant. GA treatment increased the height of wildtype and *coi1-4x* plants by 46% and 74%, respectively (Figure 13C). It did not, however, rescue the developmental delay of the third leaf in the mutant. Three weeks after MeJA treatment, wildtype plants grew out of the juvenile stage and restored their green leaves, but the *coi1-4x* failed to transition from the juvenile to adult growth phase (Figure 13B). The plant height of wildtype and *coi1-4x* treated with MeJA for three weeks showed a 41% and 55% decrease, respectively (Figure 13C, D), relative to the corresponding mock-treated control plants.

**Figure 13.**
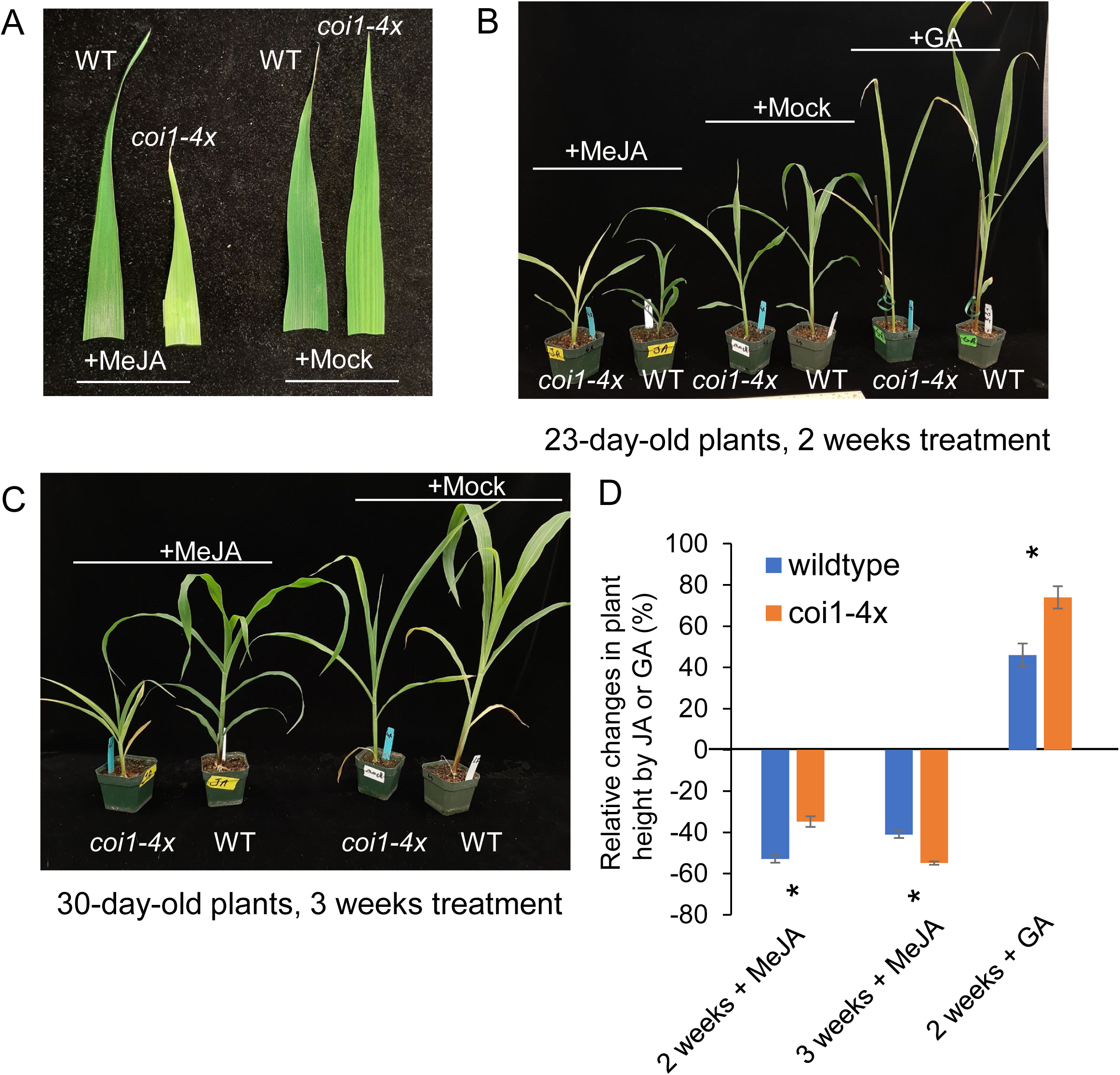
Effect of exogenous methyl jasmonate (MeJA) and gibberellic acid (GA) on wildtype and *coi1-4x* growth. (A) Leaf discoloration symptoms at 30 days post-germination and after three weeks of JA treatment. (B) Plant heights at 23 days post-germination, with mock, MeJA, or GA treatments (C) Plant heights at 30 days post-germination, with or without MeJA treatment. WT = wildtype. (D) Bar chart showing percent change in height relative to the mock-treated controls for the MeJA and GA treatments shown in panels B and C. N = 4, mean +/-s.e., two-tailed Student’s *t*-test, \**P* < 0.05.

**Figure 14.**
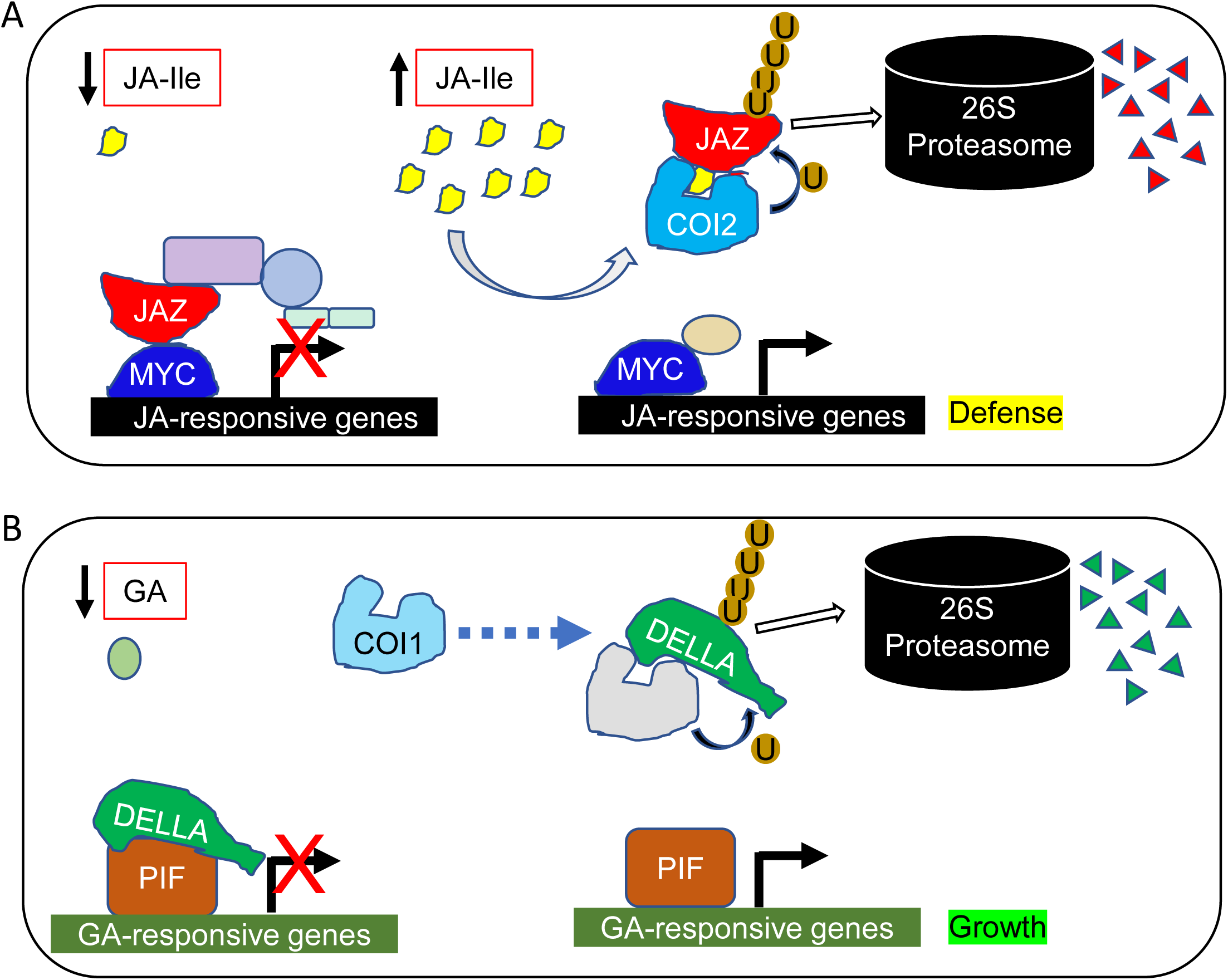
(A) A schematic model shows how maize COI2 proteins may have the classical JAZ degradation function of Arabidopsis and tomato COIs. (B) Maize COI1 proteins, on the other hand, might be F-box domain proteins that, upon binding to GA or JA, directly or indirectly degrade DELLA protein.

## Discussion

The shortened internode length and reduced plant height exhibited by the *coi1-4x* mutant are reminiscent of the constitutively-active dominant DELLA mutants or GA suppression mutants, which have been characterized in Arabidopsis (Strader et al., 2004), rice (Ueguchi-Tanaka et al., 2008), wheat (Gale and Marshall, 1973) and maize (Lawit et al., 2010; Cassani et al., 2009; Fujioka et al., 1988; Paciorek et al., 2022). The increased DELLA abundance in *coi1-4x* relative to wildtype (Figure 10) and the degradation of DELLA when co-expressed with COI in *N. benthamiana* (Figure 11) confirmed that the underlying cause of the growth deficiency in the *coi1-4x* mutant is likely increased DELLA stability. Several studies have demonstrated crosstalk between DELLA and plant defense responses (Machado et al., 2017; Hou et al., 2010; Wild et al., 2012; de Vleesschauwer et al., 2016; Qi et al., 2014; Dong and Hudson, 2022; Navarro et al., 2008; Yang et al., 2012). In Arabidopsis and rice, the JAZ repressors of the JA signaling pathway bind to and entrap DELLA to hinder its repression of the growth-related genes (Hou et al., 2010; Yang et al., 2012). As a result of JAZ stability and DELLA entrapment in the Arabidopsis *coi* and the rice *coi1a coi1b* mutants, these plants are hypersensitive to GA and grow taller than corresponding wildtype plants (Xu et al., 2002; Machado et al., 2017; Yang et al., 2012). Thus, the increased DELLA abundance and decreased growth in the maize *coi1-4x* mutant is a notable contrast to the corresponding Arabidopsis and rice *coi* knockout mutants, which are taller than the corresponding wildtype plants.

One possible explanation for the short stature of *coi1-4x* is that maize COI1 proteins have lost affinity for JAZ but not for the rest of the E3-ligase complex, thereby functioning as competitive inhibitors of COI2. The resulting enrichment of COI2 in the E3-ligase complexes, which results from COI1 depletion in *coi1-4x*, may lead to JA hypersensitivity, increased stabilization of DELLA proteins, and growth inhibition. However, expression of most of the canonical JA-responsive genes, including those encoding benzoxazinoid and terpene biosynthesis enzymes (Figures S12 and S13; Tables S7 and S8), was not significantly changed in the *coi1-4x* mutant relative to wildtype. This lack of a significant defense response at the gene expression level was in accordance with a lack of improved caterpillar growth on the *coi1-4x* mutant (Figure 4B, C).

As an alternate hypothesis, we propose that maize COI1 proteins may have acquired a new function to help maintain growth during the COI2-mediated canonical JA signaling and its associated growth penalty. This moonlighting function could form a SCF^coi1^ complex that targets DELLA instead of JAZ for its polyubiquitylation and degradation during plant defense responses. Or, perhaps maize COI1 proteins lead to the activation of some other ubiquitin ligase that triggers DELLA degradation. DELLA-RFP disappeared from the nuclei when co-expressed with COI1a, COI1c, and even COI2a in *N. benthamiana* leaves. The proteasome inhibitor BTZ restored DELLA in 60% of the nuclei, consistent with the hypothesis that maize COI proteins directly or indirectly target DELLA for degradation by the proteasome. Almost two thirds of the downregulated genes in the *coi1-4x* mutant are involved in photosynthesis and carbohydrate and cell wall metabolism (Figure 8 and Table S4). These are also among the main groups of genes that are induced by the GA signaling pathway (Ranwala and Miller, 2008; Falcioni et al., 2018; Chen et al., 2020).

DELLA is at the nexus of several signaling pathways, not limited to the crosstalk between GA and JA. In Arabidopsis, ABA inhibits GA-mediated seed germination, and overaccumulation of DELLA upregulates ABA-mediated responses. This antagonistic crosstalk was shown to be mediated through the formation of a module between the DELLA protein RGL2 and one of the three NF-YC3, 4, or 9 homologs, which subsequently binds to the promoter of the *ABI5* transcription factor and modulates the ABA-responsive genes, regardless of the ABA level (Liu et al., 2016b). In our data, one prominent group of genes that are more highly expressed in the *coi1-4x* mutant relative to all other genotypes consisted of ABA-regulated and drought-responsive genes (Figure 9 and Table S5), which is consistent with over-accumulated DELLA (Figure 10) upregulating the ABA pathway in the *coi1-4x* mutant. Downregulation of *NF-YC4* in the *coi1-4x* mutant relative to all other genotypes (Figure 8 and Table S4), on the other hand, may indicate a compensatory or regulatory feedback reaction to alleviate the effects of the DELLA-mediated ABA induction.

Both GA and JA signaling pathways have crosstalk with the auxin pathway. The interactive modules between DELLA on one hand, and ARF activators and AUX-IAA repressors, on the other hand, play crucial roles in fruit ripening and vascular development (Hu et al., 2018, 2022). Moreover, JA and auxin signaling complexes are structurally similar (Pérez and Goossens, 2013; Tal et al., 2020). Cullin1 is a scaffolding module and a shared component of the SCF ubiquitin ligase complexes involved in mediating responses to auxin and JA (Ren et al., 2005). An overabundance of Cullin1 derived from the depletion of the SCF^coi1^ complex from the four COI proteins may lead to an increased formation and activity of the SCF^TIR1^, the auxin-inducible SCF ubiquitin ligase complex (Gray et al., 2001), and hypersensitivity to auxin, assuming that the unique components of the auxin-inducible complex are not limiting. It is known that auxin inhibits seed germination in an ABA-dependent manner (Liu et al., 2013). It remains to be determined whether the auxin-related symptoms in the *coi1-4x* mutant are a consequence of auxin or other hormonal imbalances.

Our results show that 72% of the downregulated and 29% of the upregulated genes in the *coi1-4x* mutant encode proteins related to C_4_ metabolism, which exhibit bundle sheath or mesophyll-specific expression (Tables S6 and S10) (Li et al., 2010). C_4_ species have evolved more than 60 times independently in the plant kingdom in response to selective pressures from the environment (Zhou et al., 2018; Blätke and Bräutigam, 2019; Christin et al., 2013). We propose that in maize, and perhaps other C_4_ species, COI-mediated DELLA degradation might be part of the plant adaptation strategies to maintain growth in response to biotic stresses and severe environmental conditions (*e.g.*, drought, high light, and elevated temperatures), which are the driving forces in C_4_ evolution. It has long been speculated that C_4_ characteristics already existed in C_3_ plants and were only modified in C_4_ species in response to their environmental needs (Burgess et al., 2016; Wasilewska-Dębowska et al., 2022; Hibberd and Quick, 2002). A recent study showed that bundle sheath cells of rice and Arabidopsis are conditioned to synthesize proteins involved in water transport, sulfur assimilation, and JA synthesis (Hua et al., 2021).

Most downregulated genes in the *coi1-4x,* which are GA-inducible and growth-related, are also C_4_ genes with cell-specific-expression patterns (Figure 8 and Table S4). These genes encode bundle sheath-abundant photosynthetic proteins, including NadhU, which is an essential part of the NAD(P)H dehydrogenase (NDH) complex that performs cyclic electron transport (CET) in chloroplasts (Yamamoto et al., 2011). NDH levels increased from 4% of the total photosystem I (PSI) in C_3_ plants to 40% in the bundle sheath chloroplast of C_4_ to provide the high demand of ATP in the bundle sheath chloroplasts of NADP-ME C4 plants, including maize, sorghum, and *Flaveria* C_4_ species (Takabayashi et al., 2005; Wasilewska-Dębowska et al., 2022). Studies on the genus *Flaveria,* which includes both C_3_, C_4_ and intermediate species, showed that this protein complex increases up to fourteen times in the bundle sheath chloroplasts of the C_4_ *Flaveria* species compared to the C_3_ species (Nakamura et al., 2013). The NDH cyclic electron transport also protects plastoquinone, the electron receiver of cyclic electron transport, and photosystem I against excessive reduction and high light-derived reactive oxygen species, which can cause photoinhibition.

Interestingly, fibrillin 4, another photosynthetic protein that is downregulated in the *coi1-4x*, is a photosystem II (PSII) and light harvesting complex II (LHCII) associated member (Singh et al., 2010) of the fibrillin family, which plays essential roles in the biosynthesis of plastoquinone, light acclimation, and sulfur metabolism (Kim et al., 2015; Lee et al., 2020). Phosphorylation of PSII and LHCII play essential roles in the acclimation of photosynthesis to high light intensity, and this role becomes more essential in maize (Drozak and Romanowska, 2006) and other C_4_ plants (Reviewed in (Wasilewska-Dębowska et al., 2022)). Upregulation of genes coding for kinases, phosphatases, and phosphate transporters, including the sulfur transporter 3:4 (SULTR 3:4), a protein involved in the transport of sulfur (Takahashi et al., 1997) and phosphorus (Ding et al., 2020), in the *coi1-4x* mutant (Figure 9 and Table S5), might be a compensatory response. Phosphorus deficiency in the leaves of the 20-day-old *coi1-4x* seedlings (Figure 6C), strengthens this hypothesis.

In conclusion, we propose that a non-classical crosstalk between the JA and GA pathways, which is disrupted in the *coi1-4x* mutant, plays a role in the evolution of maize and perhaps other C_4_ plants. By regulating growth responses, maize COI1 proteins compensate for the canonical growth penalty associated with JA-Ile induction of plant defense responses. Future research will define the exact mechanisms of this compensatory response.

## Materials and Methods

### Plant growth conditions

Seedlings were grown in Conviron (Winnipeg, Canada) growth chambers under a light intensity of 500 μmol/m^2^/sec, a 16:8 h light:dark cycle, 25 °C light, 22 °C dark, and 50% relative humidity, and in a soil mix containing 35% peat moss, 10% vermiculite, 35% baked clay, 10% sand, 10% sterilized topsoil. Methyl Jasmonate (MeJA) and gibberellic acid (GA) treatments were done on plants that were germinated in soil or germination pouches. In the latter case, seeds were germinated in CYG germination pouches (Mega-International.com).

### Phylogenetic tree and protein alignment

The accession numbers and sequences for the maize six COI proteins were obtained from the maizeGDB (https://www.maizegdb.org/). Blastp (protein-protein BLAST; https://blast.ncbi.nlm.nih.gov/) was used for pairwise alignments and calculating the percentage identity between COI proteins. The paralogous COI sequences from other organisms were obtained from Phytozome (https://phytozome-next.jgi.doe.gov/) and were aligned with the maize proteins using multAlin (http://multalin.toulouse.inra.fr/multalin/). A phylogenetic tree of COI proteins from different species was created using phylogeny.fr (http://www.phylogeny.fr).

### Transposon insertion lines and genotyping

The *coi2 Mu* transposon insertion alleles were identified from the UniformMu Transposon Resource (https://curation.maizegdb.org/documentation/uniformmu/index.php) (Settles et al., 2007). The *coi1a, coi1b,* and *coi1c* mutants were created by remobilizing nearby *Ds* transposon insertions. *Ds* transposon insertions that were tightly linked to maize *COI1a, COI1b,* and *COI1c* genes (I.S07.1293, I.W06.0524B, and I.S07.1819, respectively) were identified through the MaizeGDB webpage (https://www.maizegdb.org/). The identities of these lines were verified with primers designed to specifically amplify a *Ds* junction fragment. Verified lines were crossed with *Activator* (*Ac*) to induce *Ds* transposition. F1 seeds from these crosses were planted and screened within two weeks after germination using primers designed to the *Ds* end sequence and the *COI* sequence of interest (Table S11). A total of 4,004 (*COI1a*), 6,160 (*COI1b*), and 1,800 (*COI1c*) maize seedlings were screened by PCR to find transposon insertions in each *COI* gene. Amplification fragments indicated a potential insertion in a *COI* gene, and genotypes were verified through successive PCR validation screens using different primer sets that amplified *Ds* and *COI*, followed by Sanger sequencing to identify the insertion site (Table S3). Seedlings carrying novel *COI::Ds* alleles were transplanted and grown to maturity in a greenhouse and self-pollinated to recover progeny. Seeds from these *coi Ds* insertion lines were harvested and planted for screening of homozygous *Ds* insertions. Homozygous *coi Ds* insertion lines were then further characterized in this study. Detailed methodology of *Ac*/*Ds* tagging has been described previously, and the *coi1d* mutant was identified from this maize *Ds* transposon insertion collection (Vollbrecht et al., 2010; Ahern et al., 2009). Genotyping of the mutant and wildtype alleles of each *COI* gene was performed using the PCR primer pairs listed in Table S11.

### Insect bioassays

Eggs of *S. frugiperda* and *S. exigua* were purchased from Benzon Research (www.benzonresearch.com) and were hatched at 28 °C on fall armyworm diet and beet armyworm diet (Southland Products, www.southlandproducts.net), respectively. Neonate larvae were confined on three-week-old maize plants using micro-perforated plastic bread bags (www.amazon.com) that were sealed around the base of the plants with wire twist ties. After ten days, caterpillars were harvested and weighed.

### COI and DELLA construct preparation, plant infiltration, and imaging

To prepare the BiFC assay, the coding regions of *COIA1*, *COI1c*, and *COI2a* (W22 accession numbers in Table S1), 15 *JAZ* genes (W22 accession numbers in Table S9), and the *DWARF9* (*DELLA*) gene (accession number: Zm00004b011408), were either amplified by PCR or synthesized by Twist Bioscience (https://www.twistbioscience.com/) and cloned by restriction cloning using PacI and SpeI or AscI into the BiFC vectors p2YC and p2YN (Kong et al., 2014) to generate COI-cYFP, and JAZ-nYFP, respectively. The primers used for vector construction are listed in Table S11.

To prepare constructs for the subcellular localization, the open reading frames of *COI1a, COI1c*, *COI2a,* and *DELLA* (*DWARF9*) were amplified and cloned into pDONR 207 using BP clonase recombination (Invitrogen, Carlsbad, CA, USA), before being transferred with a second recombination reaction (LR, Invitrogen) into the vector pEAQ-EGgW (Berthold et al., 2019). The primers used for vector construction are listed in Table S11. The recombinant vectors were transformed into *Agrobacterium tumefaciens* strain GV3101 and grown in LB medium with 50 mg l^−1^ kanamycin and 50 mg l^−1^ rifampicin for 1 d. After centrifugation for 10 min at 2000 x *g*, the bacteria were collected and resuspended in an infection solution (10 mM MES, 10 mM MgCl_2_ and 200 μM acetosyringone). The prepared suspensions (A600 nm = 0.5) were infiltrated into young but fully expanded *N. benthamiana* leaves using a needleless syringe. Two to three days later, yellow fluorescent protein (YFP) (for BIFC) and enhanced green fluorescent protein (EGFP) for subcellular localization were monitored using a laser confocal scanning microscope (LSM 800, Zeiss).

### Expression analysis by RNA sequencing

Tissue was collected from the sixth leaf, 20 days after germination, at the end of the light period. Leaves were sprayed with either 0.02% MeJA or solvent-only solutions for 12 hours, with one-hour intervals. Roughly 100 mg of leaf tissue from the middle of the leaf without the midrib was placed into collection tubes. RNA extraction and 3’ RNA sequencing were performed at the Cornell Institute of Biotechnology Genomic Facility (https://www.biotech.cornell.edu/core-facilities-brc/facilities/genomics-facility), as described previously (Kremling et al., 2018)

Single-end Illumina reads generated from 3’ RNA-Seq libraries were processed to remove adaptor and low-quality sequences using Trimmomatic (v0.38) (Bolger et al., 2014) and to trim polyA/T tails using PRINSEQ++ (Cantu et al., 2019). The resulting reads were aligned to the SILVA rRNA database (Quast et al., 2013) using Bowtie (Langmead et al., 2009) to remove ribosomal RNA contamination. The cleaned high-quality RNA-Seq reads were aligned to the maize W22 reference genome (Springer et al., 2018) using HISAT2 (v2.1) (Kim et al., 2019) with default parameters. Based on these alignments, raw read counts were derived for each gene and normalized to reads per million mapped reads (RPM). Differential gene expression analyses were performed using DESeq2 (Love et al., 2014).

For heatmap preparations, the RPM data were transformed by log(1+RPM) prior to clustering. The pairwise comparative *p* values (<0.05) were used for the sequential sorting to make the gene lists used in heatmaps. Heatmaps were produced in R (https://www.r-project.org/) and Venn diagrams were made using FunRich (http://www.funrich.org/).

### Inductively coupled plasma atomic absorption emission spectroscopy

Elemental analysis determinations were performed using ICP-MS. Less than 200 milligrams of leaf tissue were digested with a cocktail of HNO_3_ and perchloric acid (1:1 ratio), diluted in 10 ml of 5% HNO_3_, and analyzed using Sciex Inductively coupled argon plasma (AB Sciex LLC, Framingham, MA), as described previously (Cobb et al., 2021).

### Photosynthetic measurements

Photosynthetic efficiency was assessed by measuring gas exchange at a fixed CO_2_ concentration of 400 µmol mol^-1^ (GasExA), using an LI-6800 portable photosynthesis system (Li-Cor Inc., Lincoln, NE, USA). The efficiency of the photosystem II photochemistry and other light response parameters was assessed at a fixed light intensity of 2000 μmol m^-2^s^-1^.

### Protein Extraction, and Western Blotting

Total protein was extracted from 60 mg of leaf material as described previously (Feiz et al., 2012). Briefly, the ground leaf material was mixed with 250 µl of the extraction buffer (50 mM Tris-HCl, pH 7.5, 150 mM NaCl, 1% Triton X-100, 1 mM EGTA and 1 mM dithiothreitol) with 1% protease inhibitor cocktail (04693116001, Roche) and diluted 1x with the 2x loading buffer (65.8 mM Tris-HCl, pH 6.8, 2.1% SDS, 26.3% (w/v) glycerol, 0.01% bromophenol blue). A volume corresponding to 150 µg of tissue was analyzed using 10% SDS-polyacrylamide gels. Proteins were analyzed by transfer to polyvinylidene difluoride (PVDF) membranes. Immunodetection of the maize DELLA proteins was performed using anti-SLR1 primary antibody (Cosmo Bio USA, Carlsbad, California) (2:10,000) and anti-rabbit IgG, horseradish peroxidase conjugated secondary antibody (www.promega.com) (1:10,000). The corresponding molecular weight for the band in western blotting (70 kDa) was consistent with the expected size of maize DELLA proteins (65-66 kDa). The chemiluminescence signal was detected with the Pierce^TM^ ECL Plus Western Blotting Substrate on a GelDoc Go Imaging (BIO-RAD).

Bortezomib (BTZ, Selleckchem.com) at 20µM concentration was infiltrated into the *N. benthamiana* leaves 24 hours after leaf infiltration with the plasmids expressing the DELLA-RFP, and COI1a-EGFP or COI1c-EGFP proteins. Sixteen hours after BTZ treatments, leaves were used for confocal microscopy or immunoblot analyses.

### Accession numbers

Raw RNA-Seq reads have been deposited in the NCBI BioProject database under accession PRJNA951759.

## List of Supplemental Materials

### Supplemental Figures

**Figure S1.** Amino acid identity of Arabidopsis and maize COI proteins

**Figure S2.** Alignment of *COI* genes from different plant species

**Figure S3.** Subcellular locations of the maize COI1a, COI1c, and COI2a proteins

**Figure S4.** BIFC between the maize COI1A, COI1B or COI2a and seven JAZ proteins

**Figure S5.** Independent repeat of the COI-JAZ BIFC experiment

**Figure S6**. Segregation patterns of selfed *coi2a/coi2a COI2b/coi2b* and *COI2a/coi2a coi2b/coi2b* mutants

**Figure S7**. Test crosses between *coi2a/coi2a COI2b/coi2b* and *COI2a/coi2a coi2b/coi2b* and the wildtype W22 maize

**Figure S8.** Expression of maize *COI2* genes

**Figure S9.** Proteomics data show pollen-specific abundance of maize COI2a protein.

**Figure S10.** Growth phenotypes of *coi1-4x* compared to wildtype and double mutants

**Figure S11.** Microelements, macroelements, and photosynthesis assays.

**Figure S12.** Heatmap showing expression of maize benzoxazinoid biosynthesis genes

**Figure S13.** Heatmap showing expression of the maize terpene synthase genes.

**Figure S14.** Heatmap showing expression of the maize *JAZ* genes

**Figure S15.** DELLA protein decreases and increases after treatment of the 5-day-old seedlings with GA and MeJA

**Figure S16.** Co-localization of DELLA-RFP and COI1a-EGFP

### Supplemental Tables

**Supplemental Table S1.** Cross-reference of maize *COI* gene names

**Supplemental Table S2.** COI protein sequences that are included in the phylogenetic tree in Figure 1

**Supplemental Table S3.** Flanking sequences of transposon insertion sites in maize COI gene

**Supplemental Table S4.** List of genes that are downregulated in coi1-4x

**Supplemental Table S5.** List of genes that are upregulated in coi1-4x

**Supplemental Table S6.** List of downregulated genes that are bundle sheath and mesophyll specific

**Supplemental Table S7.** Expression levels of benzoxazinoid biosynthesis genes, with and without MeJA treatment

**Supplemental Table S8.** Expression levels of terpene synthases, with and without MeJA treatment

**Supplemental Table S9.** Expression levels of maize *JAZ* genes, with and without MeJA treatment

**Supplemental Table S10.** List of upregulated genes that are bundle sheath and mesophyll specific

**Supplemental Table S11.** Primers used for genotyping and gene amplification

### Supplemental Datasets

**Supplemental Dataset S1.** RNA-Seq data for all annotated genes, pairwise comparisons

**Supplemental Dataset S2.** Expression levels of differentially expressed genes

## Acknowledgments

We thank Shree Giri at the USDA, Robert W. Holly Center for Agriculture and Health, Plant Soil and Nutrition Research for ICP atomic absorption emission spectroscopy; Filipe Aiura Namorato (USDA, Ithaca), Eric Craft (USDA, Ithaca), Eric Schmelz (UC San Diego), Brad Nelms (University of Georgia), Honghe Sun (BTI), Judith Kolkman (Cornell University), Fay Wei Li (BTI), Venkatesh Thirumalaikumar (BTI), Cynthia Holland (Williams College), Amber Hotto (BTI) for their conceptual and/or technical advice; Dong-Lei Yang and Mingzhu Wang (Nanjing Agricultural University) for kindly providing the SLR1 antibody; and Miyanko Ueguchi-Tanaka from Nagoya University for the helpful information regarding the process to purchase their SLR1 antibody #2 from Cosmo Bio USA; and Corinne Schmitt-Keichinger (Université de Strasbourg) for kindly providing the modified and tagged versions of the pEAQ vectors.

This project was supported by NSF award 2019516 and USDA award 2021-67014-342237 to GJ, an International Postdoc Scholarship from the Punjab Higher Education Commission to IG, and USDA-AFRI NIFA Postdoctoral Fellowship 2014-67012-22269 to CS.

## FIGURES

**Figure S1.**
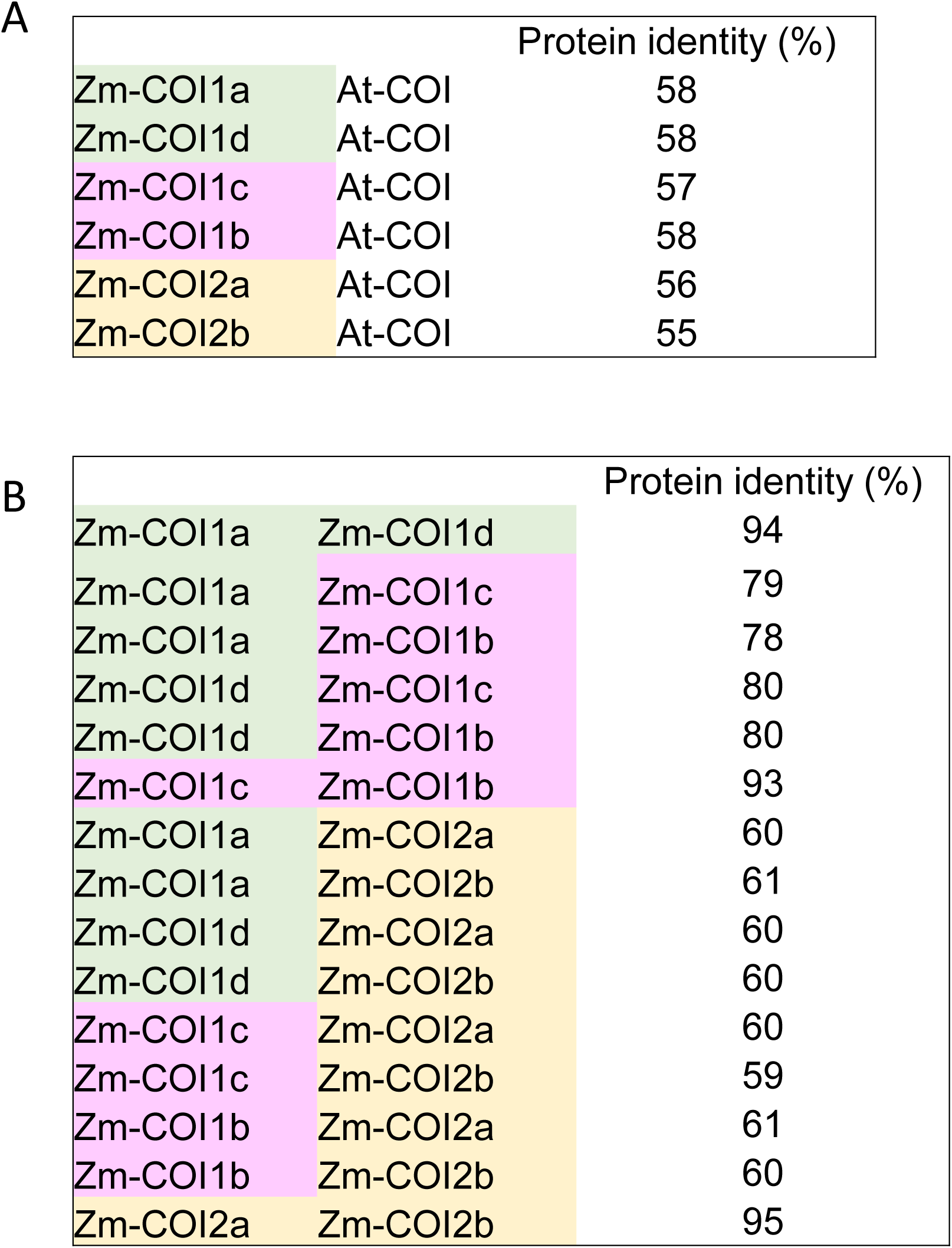
(A) Percent amino acid identity between maize COI proteins and Arabidopsis COI proteins. (B) Percent amino acid identity among the six maize COI proteins. The maize proteins are color-coded based on pairwise similarity in the phylogenetic tree in Figure 1.

**Figure. S2.**
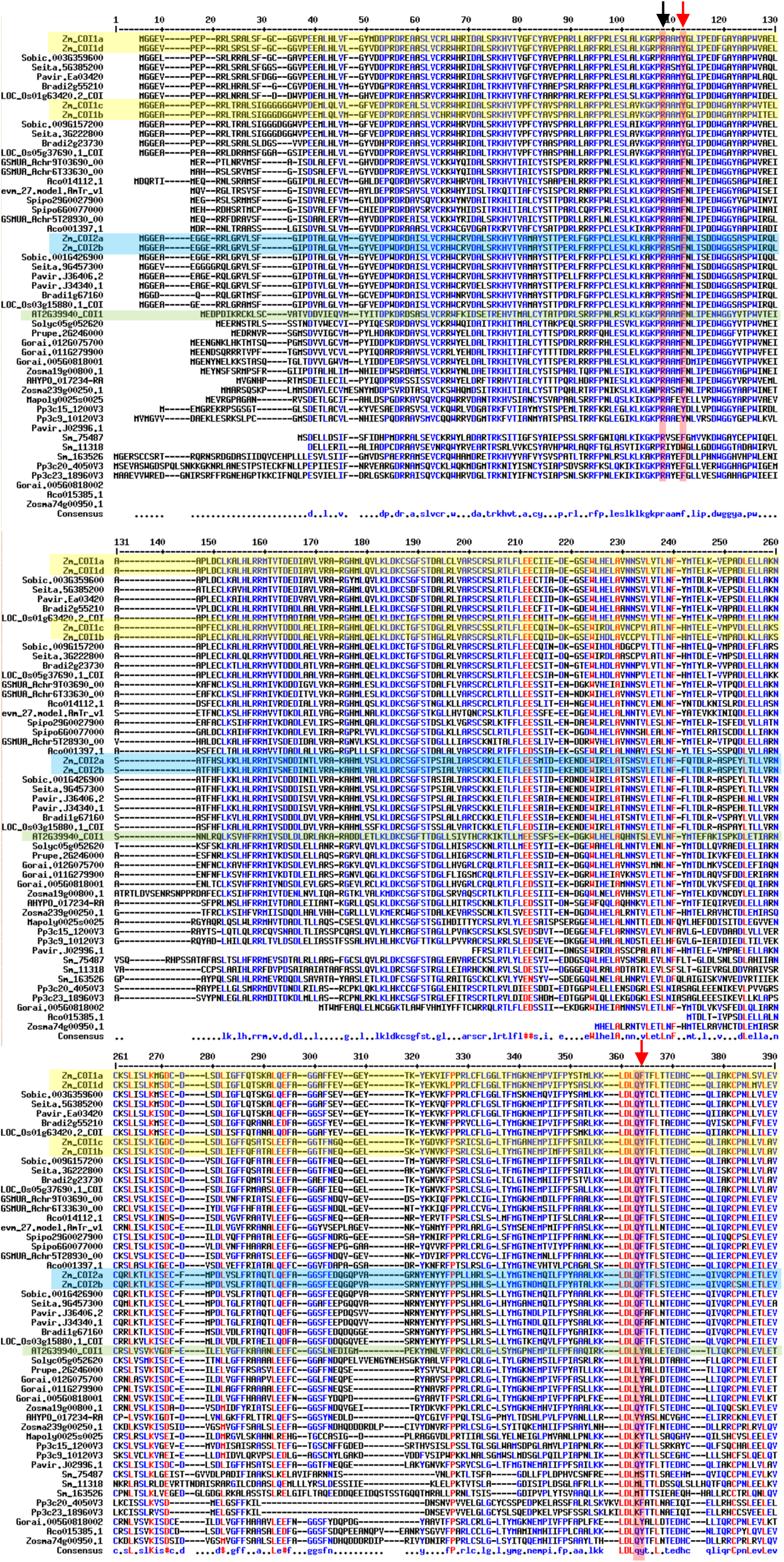
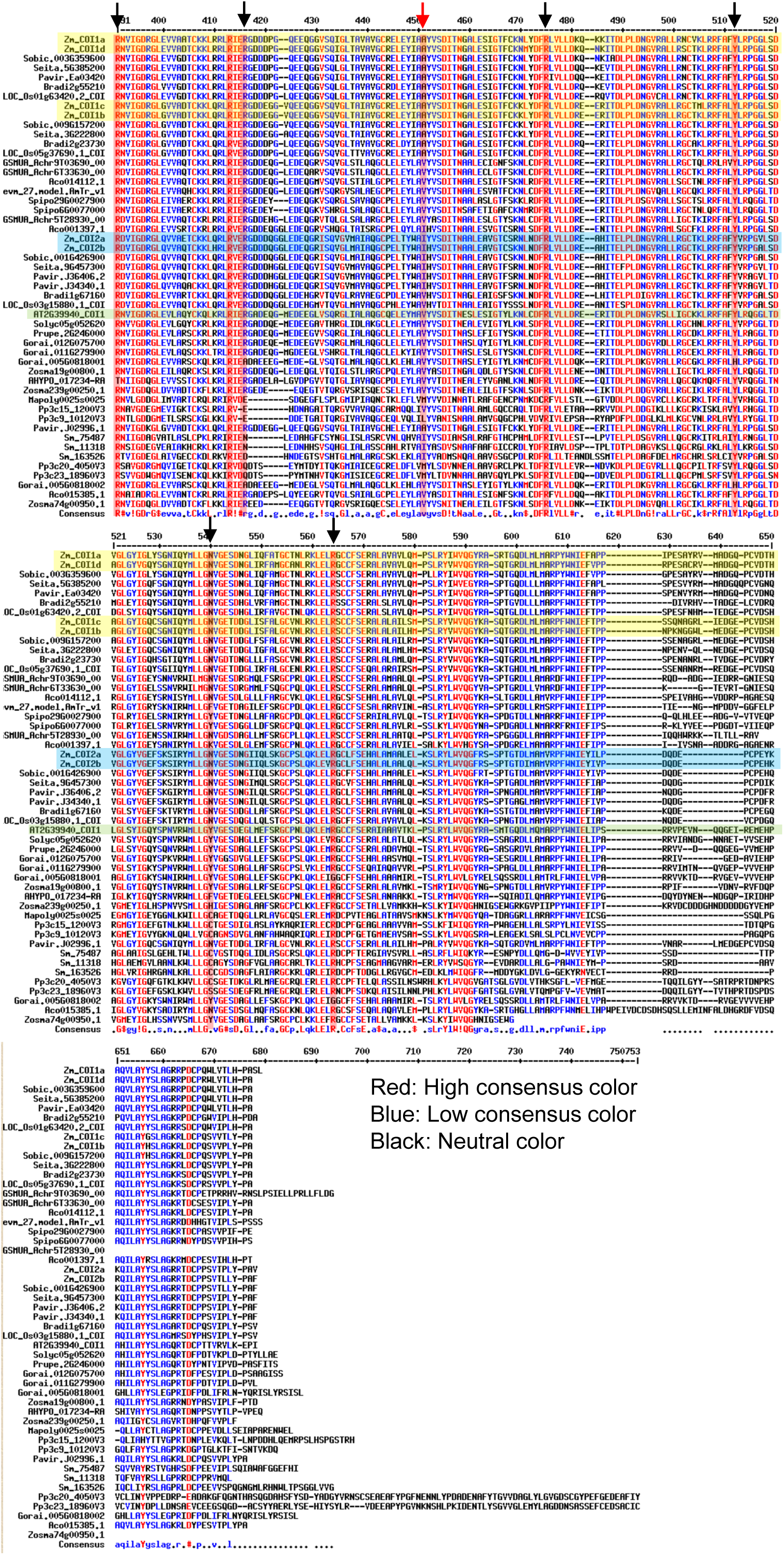
Alignment of *COI* genes from different plant species. Red arrows show the active site amino acids of the Arabidopsis COI that are modified in the COI clades across the Poaceae. Black arrows show the active site amino acids that are not modified.

**Figure S3.**
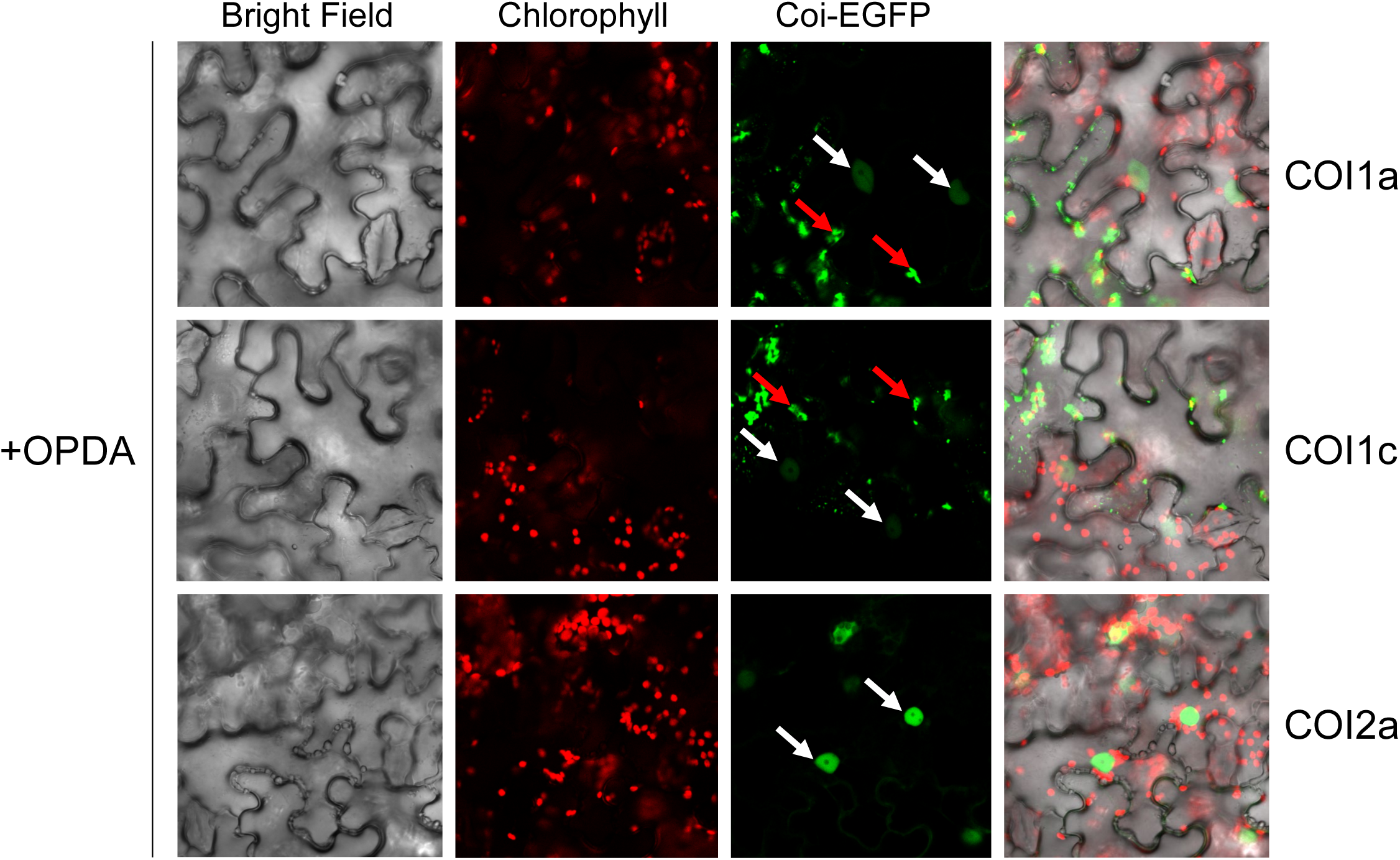
Subcellular locations of the maize COI1a, COI1c, and COI2a proteins show that COI2a is more targeted to the nuclei than the COI1s. Confocal images were taken at 24 h after infiltrating plasmids expressing the EGFP-COI fusion proteins into the *Nicotiana benthamiana*. Leaves were sprayed with 12-oxo-phytodienoic acid (OPDA), two hours before imaging. White arrows indicate the nuclei, and red arrows represent cytosolic condensates. Scale bars = 25 μm.

**Figure S4.**
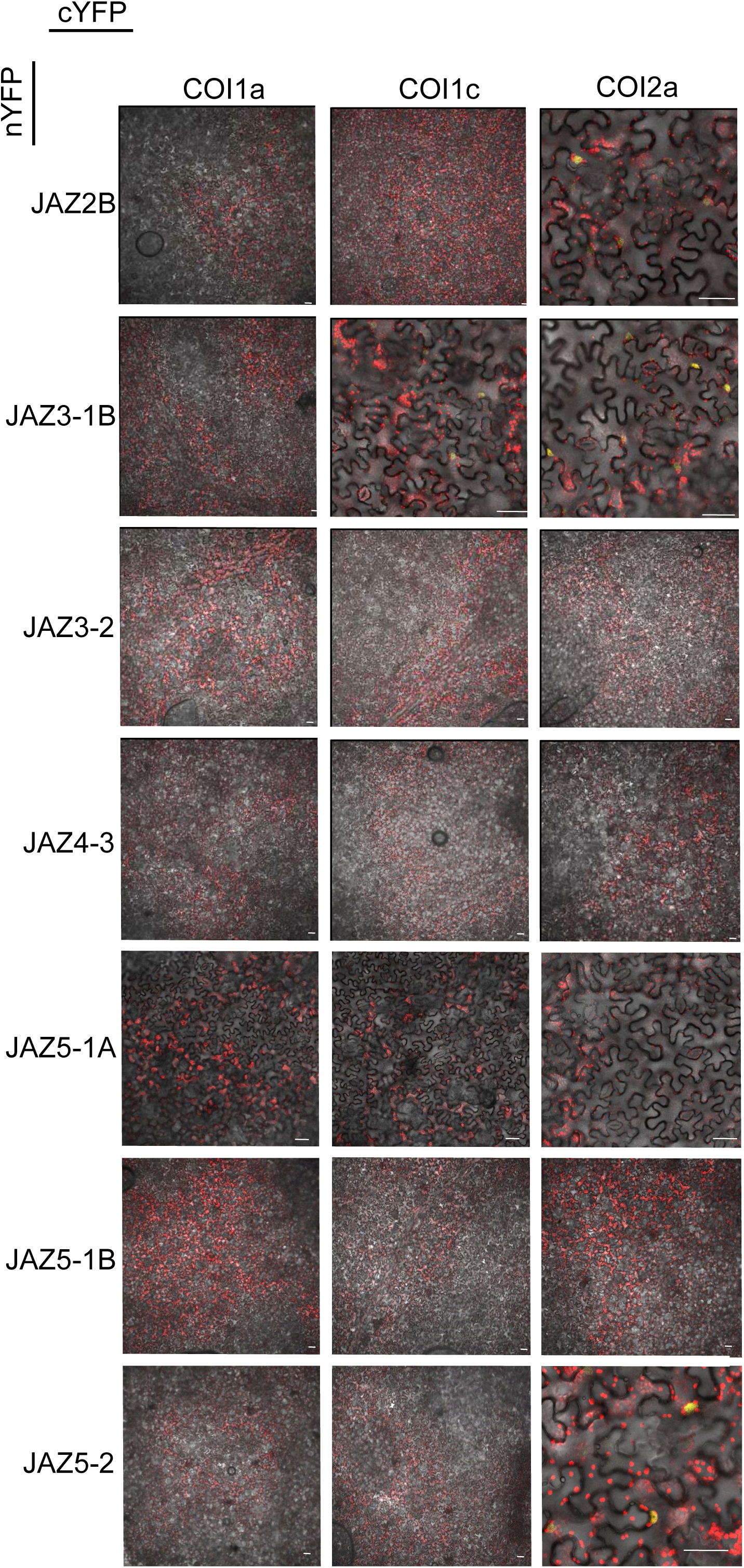
Bimolecular fluorescence complementation (BIFC) between the maize COI1a, COI1c, and COI2c and seven JAZ proteins that did not show interactions with any of the COI proteins. Scale bars = 50 μm.

**Figure S5.**
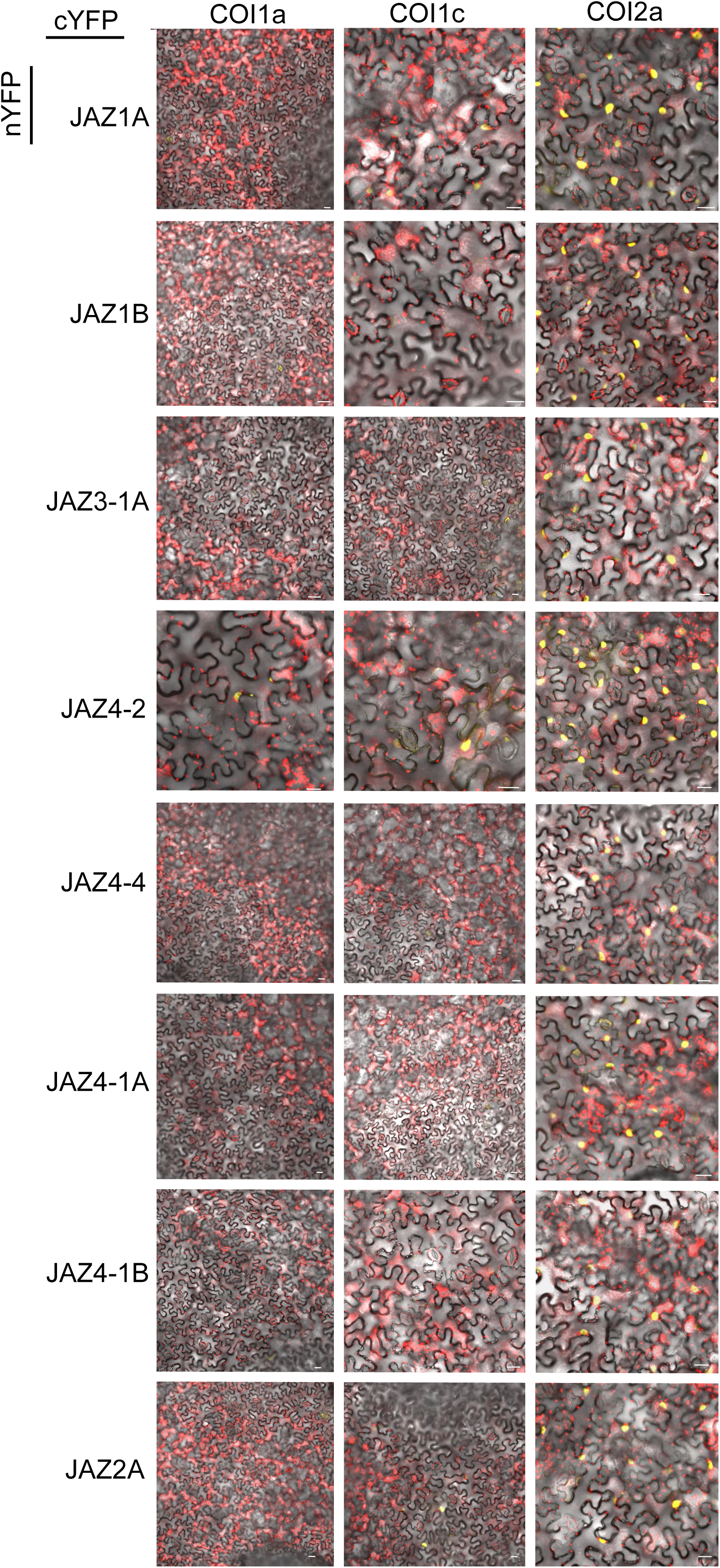
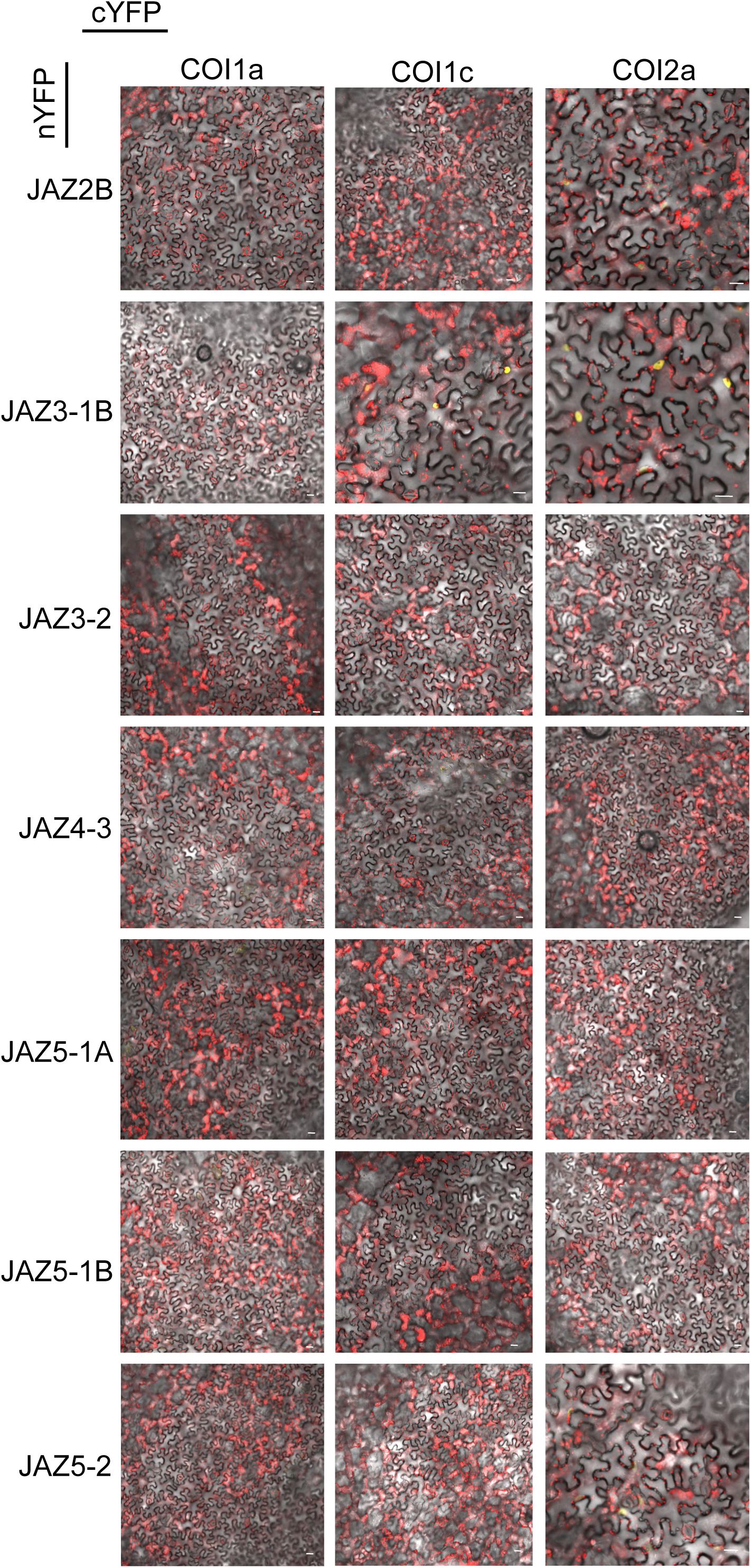
An independent repeat of the bimolecular fluorescence complementation (BIFC) experiment that is presented in Figures 3 and S3. Scale bars= 50 μm.

**Figure S6.**
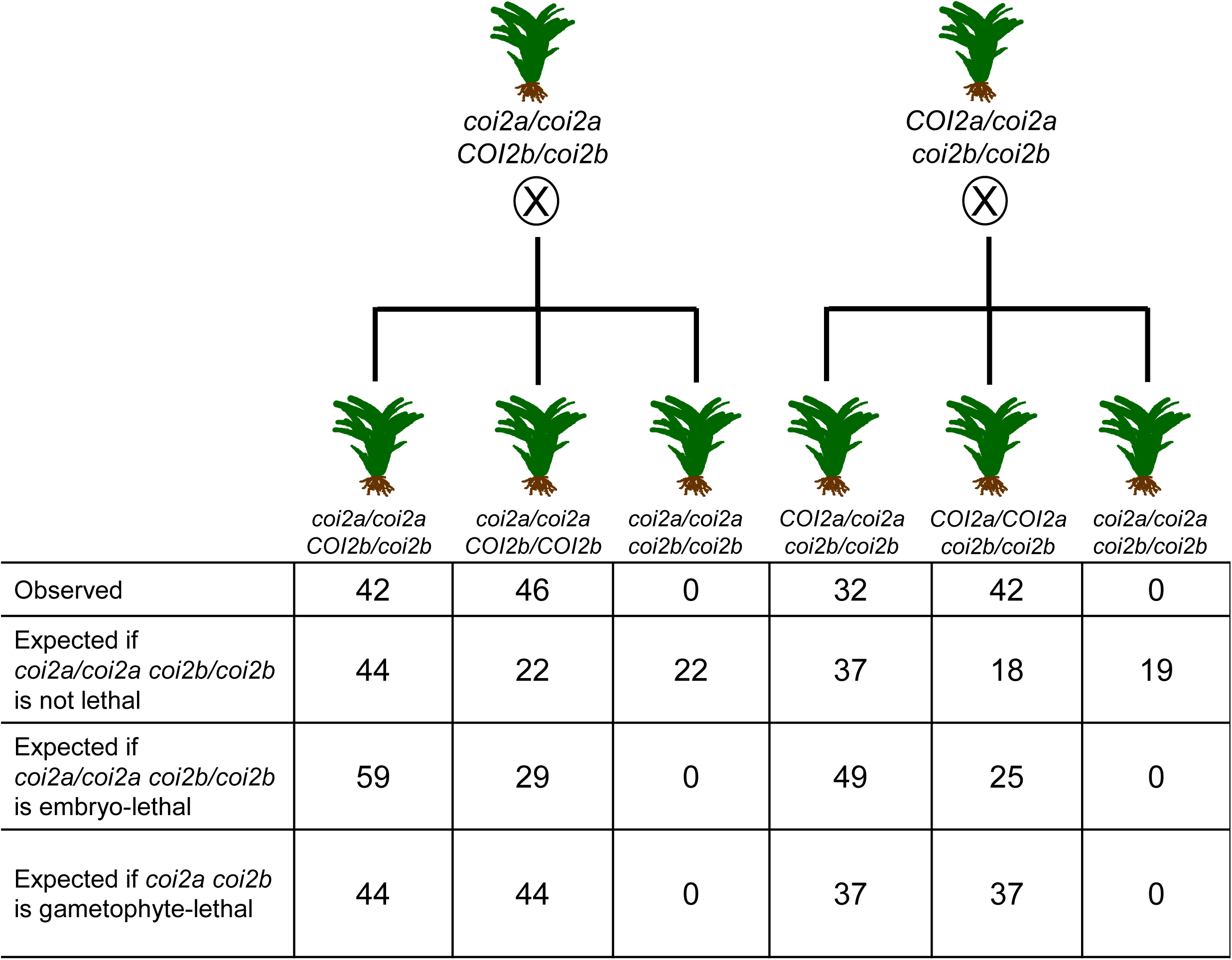
Segregation patterns of seeds derived from *coi2a/coi2a COI2b/coi2b* and *COI2a/coi2a coi2b/coi2b* plants show that there are no homozygous *coi2a/coi2a coi2b/coi2b* mutants among the progeny. This indicates that either male or female *coi2a coi2b* gametes do not survive.

**Figure S7.**
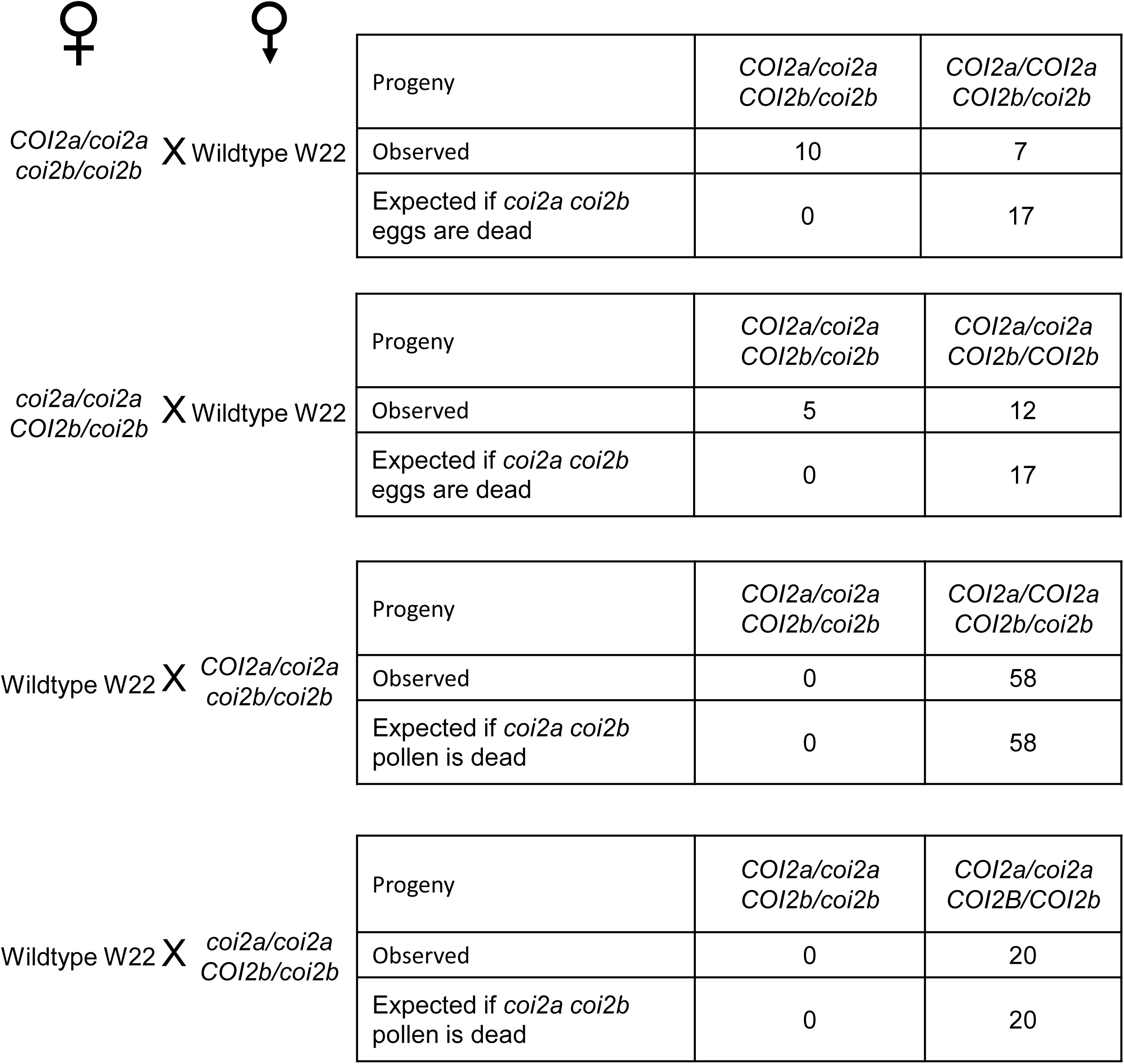
Test crosses between *COI2a/coi2a coi2b/coi2b* or *coi2a/coi2a COI2b/coi2b* and wildtype maize inbred line W22 indicate that *coi2a coi2b* pollen is not viable.

**Figure S8.**
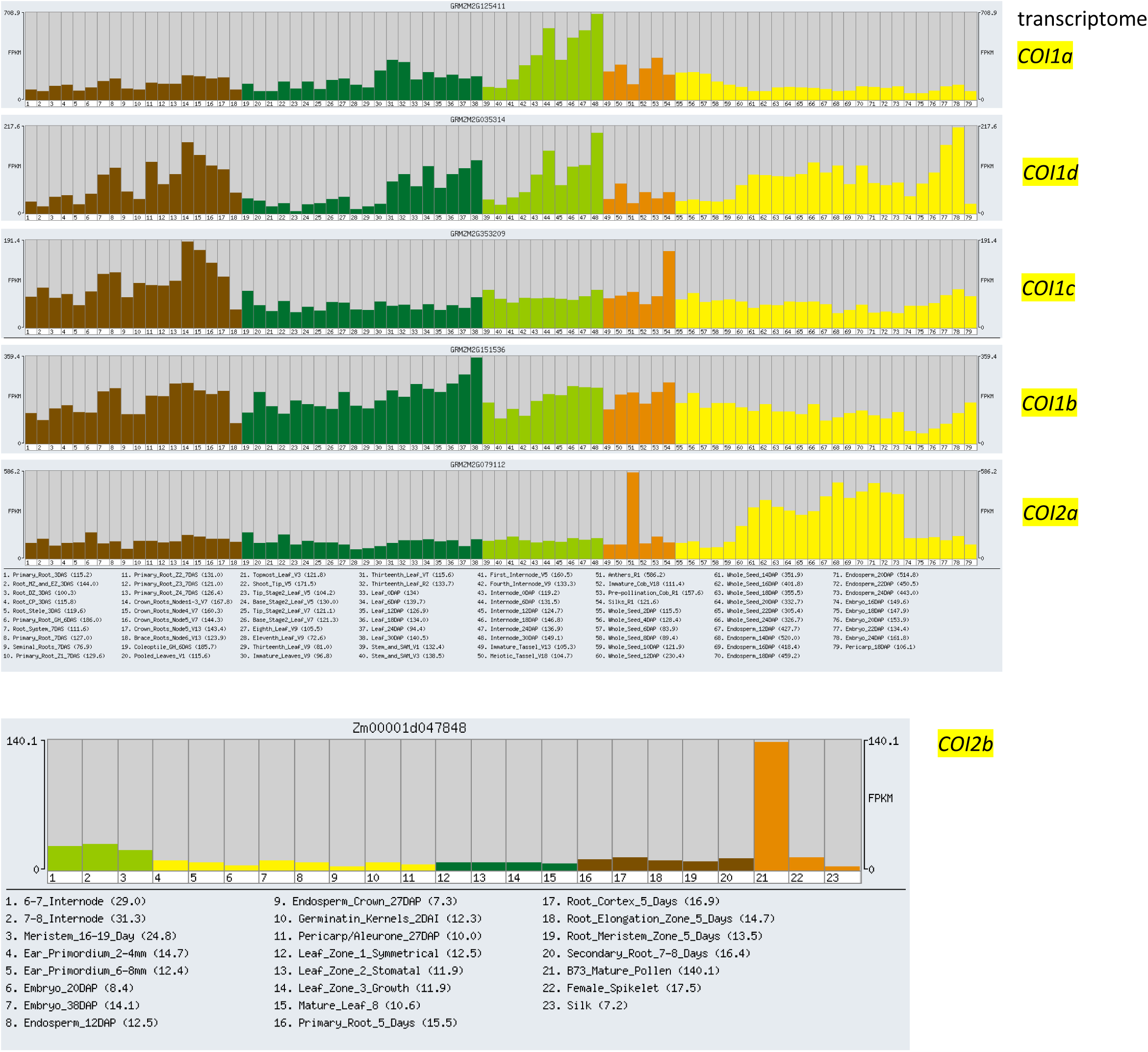
Transcriptome data show highest expression of maize *COI2* genes in anthers. Data were plotted from the Maize Genome Database (www.maizegdb.org)

**Figure S9.**
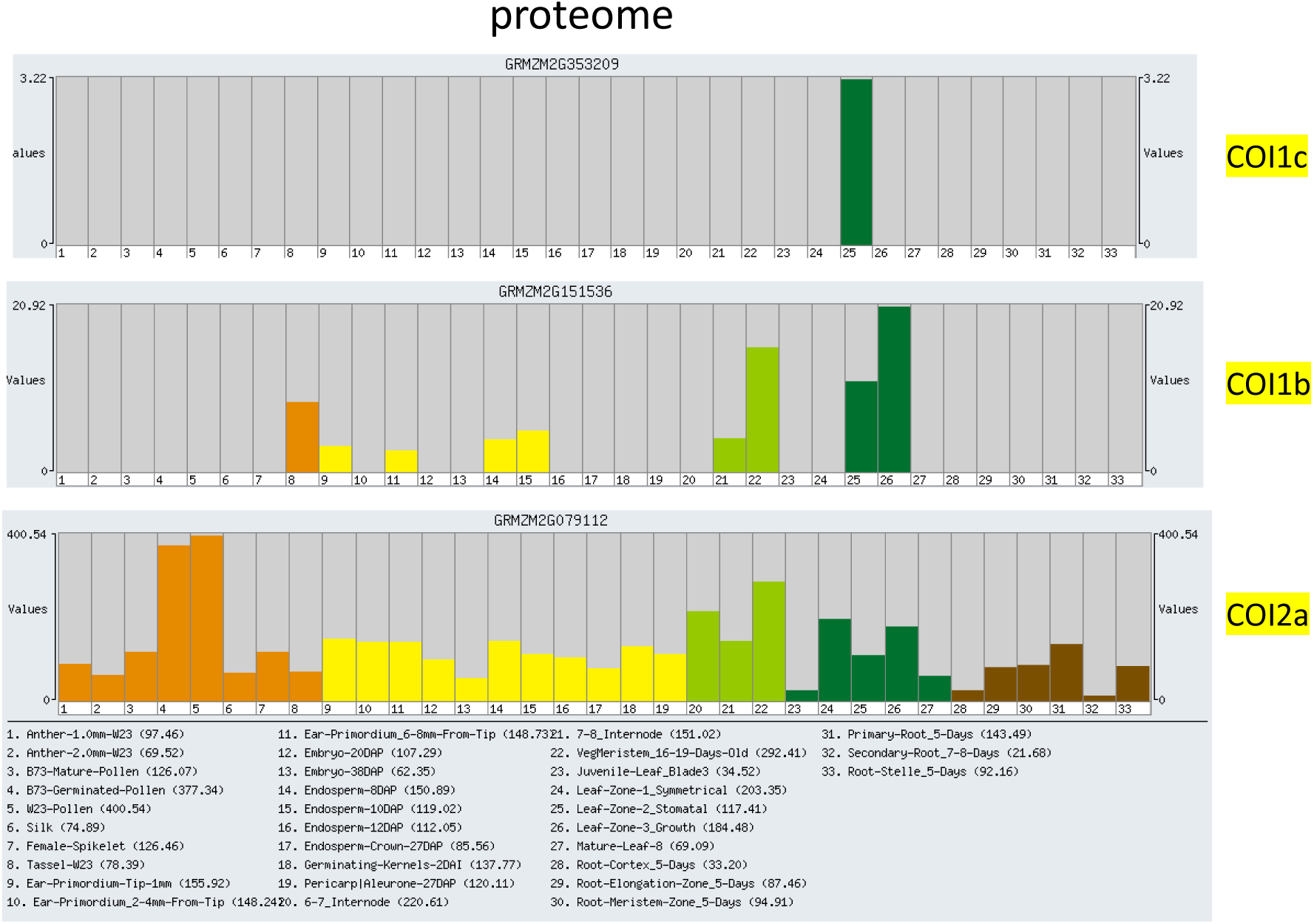
Proteomics data show pollen-specific abundance of maize COI2a protein. Data are from: Walley et al (2016), Integration of omic networks in a developmental atlas of maize, *Science*, 353:814-818,

**Figure S10.**
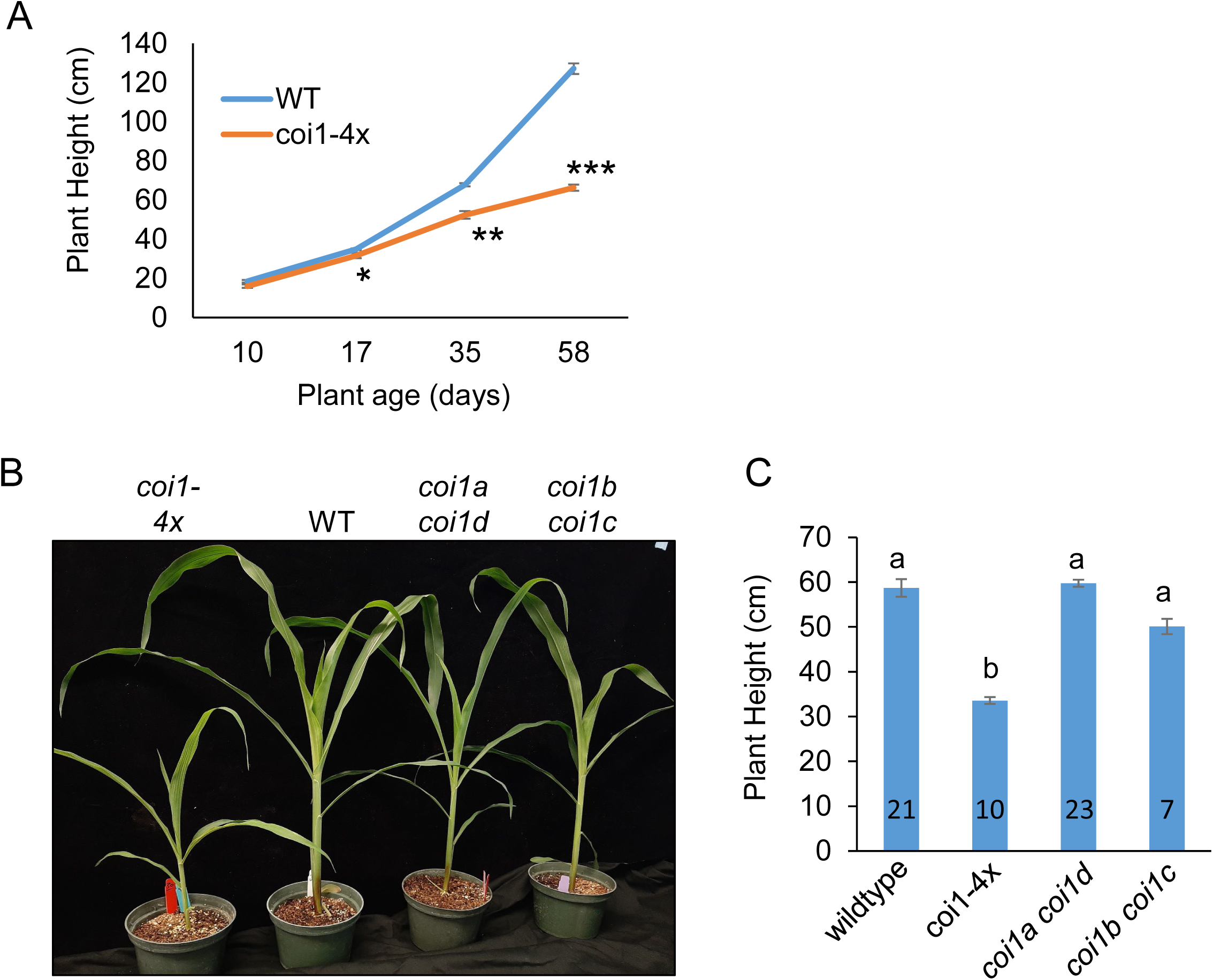
Phenotypic comparisons between the *coi1* quadruple mutant (*coi1-4x*) and the corresponding double mutants, *coi1a coi1d* and *coi1b coi1c*, and wildtype W22. (A) plant heights were compared between *coi1-4x* and wildtype at the indicated ages. N =1 0 at 10, 17, and 35 days, and N= 6 at 58 days. Student’s *t*-test, \**P* < 0.05, ** *P* < 0.01, *** *P* < 0.001. (B, C) Plant heights at 20 days after germination. Numbers in bars = sample sizes, mean +/-s.e., letters indicate differences P < 0.05 using ANOVA followed by Tukey’s HSD test.

**Figure S11.**
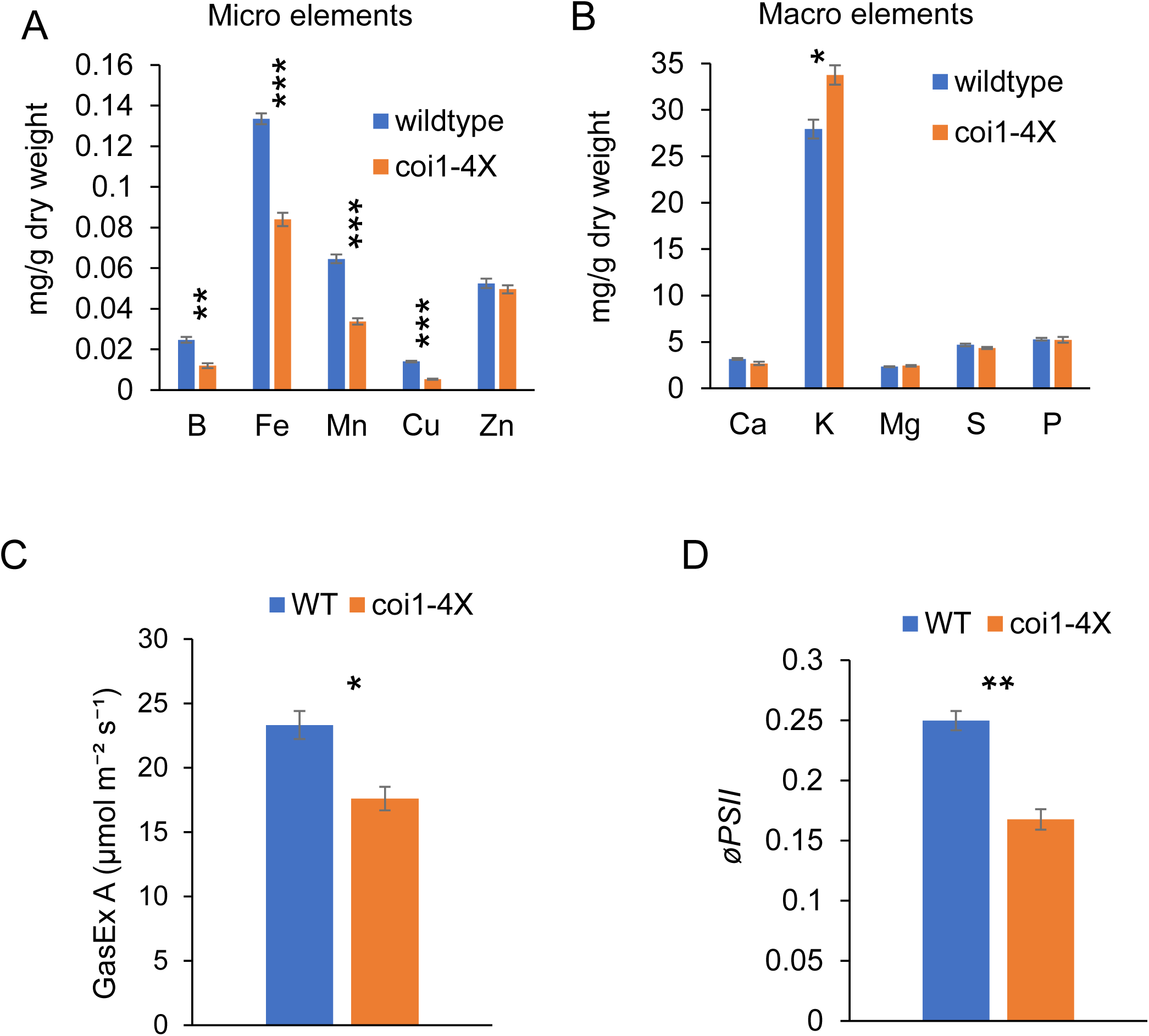
Microelements, macroelements, and photosynthesis assays. (A) Microelements and (B) macroelements in 13-day-old seedlings, N = 5, mean +/-s.e., Student’s *t*-test with (* *P*<0.05, ** *P*<0.01, *** *P*<0.001). Macro and micronutrients were measured in dried tissue of the third leave by inductively coupled plasma – atomic absorption emission spectroscopy (ICP-AES). (C) Leaf CO_2_ assimilation rate (GasEX A) at 400 µmol mol⁻¹ CO_2_ and (D) the quantum yield of the photosystem II phytochemistry (*øPSII*) at 2000 μmol m^-2^ s ^-1^ of actinic light (B) were measured at 28 days after germination, respectively. Mean ± se., n = 5, two-tailed Student’s *t*-test, \**P* < 0.05, \*\**P* < 0.01, \*\*\**P* < 0.001, WT – wildtype.

**Figure S12.**
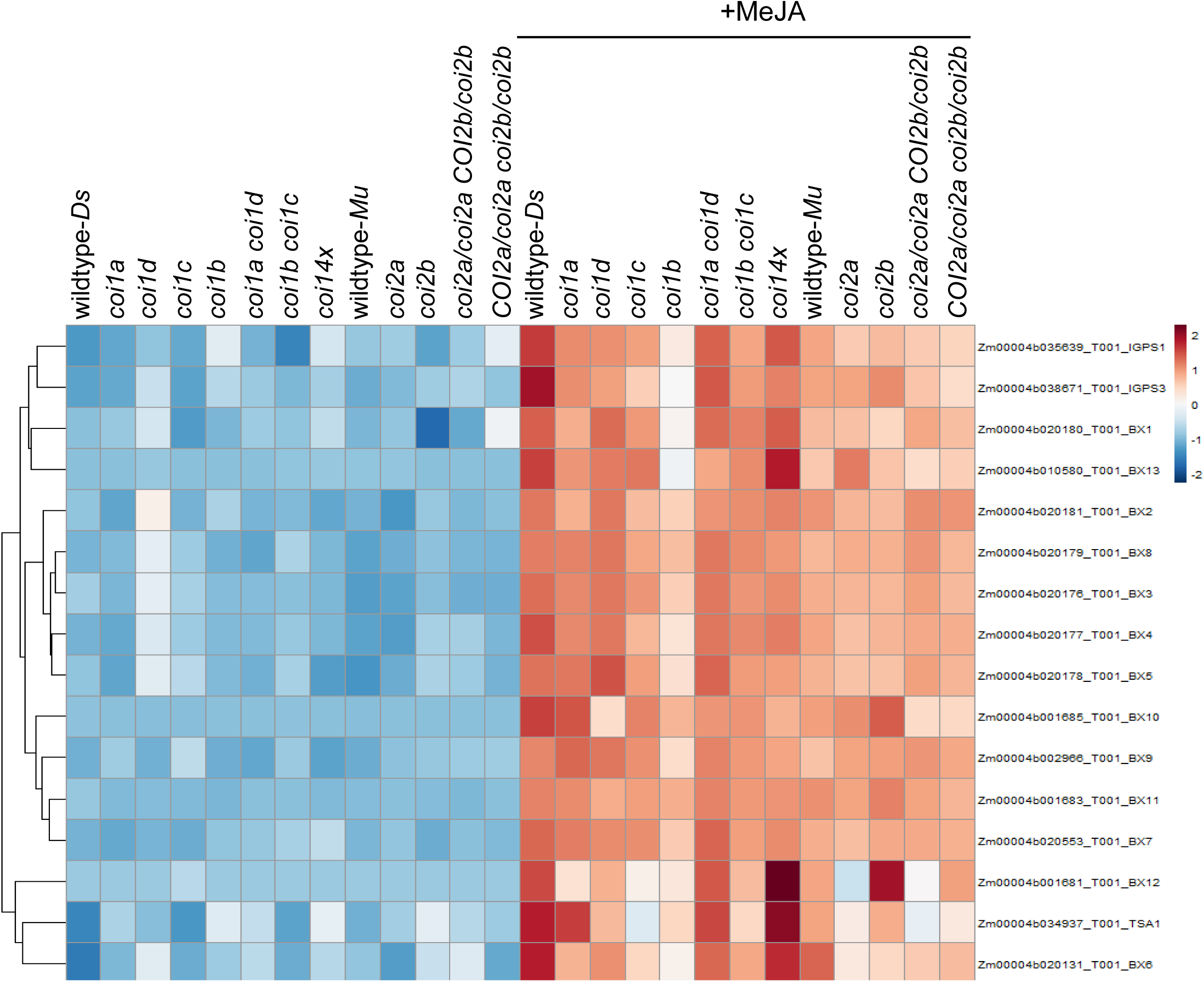
Heatmap showing expression of the benzoxazinoid biosynthesis genes with and without MeJA induction. The expression levels of the genes used in this heatmap are presented in Table S7.

**Figure S13.**
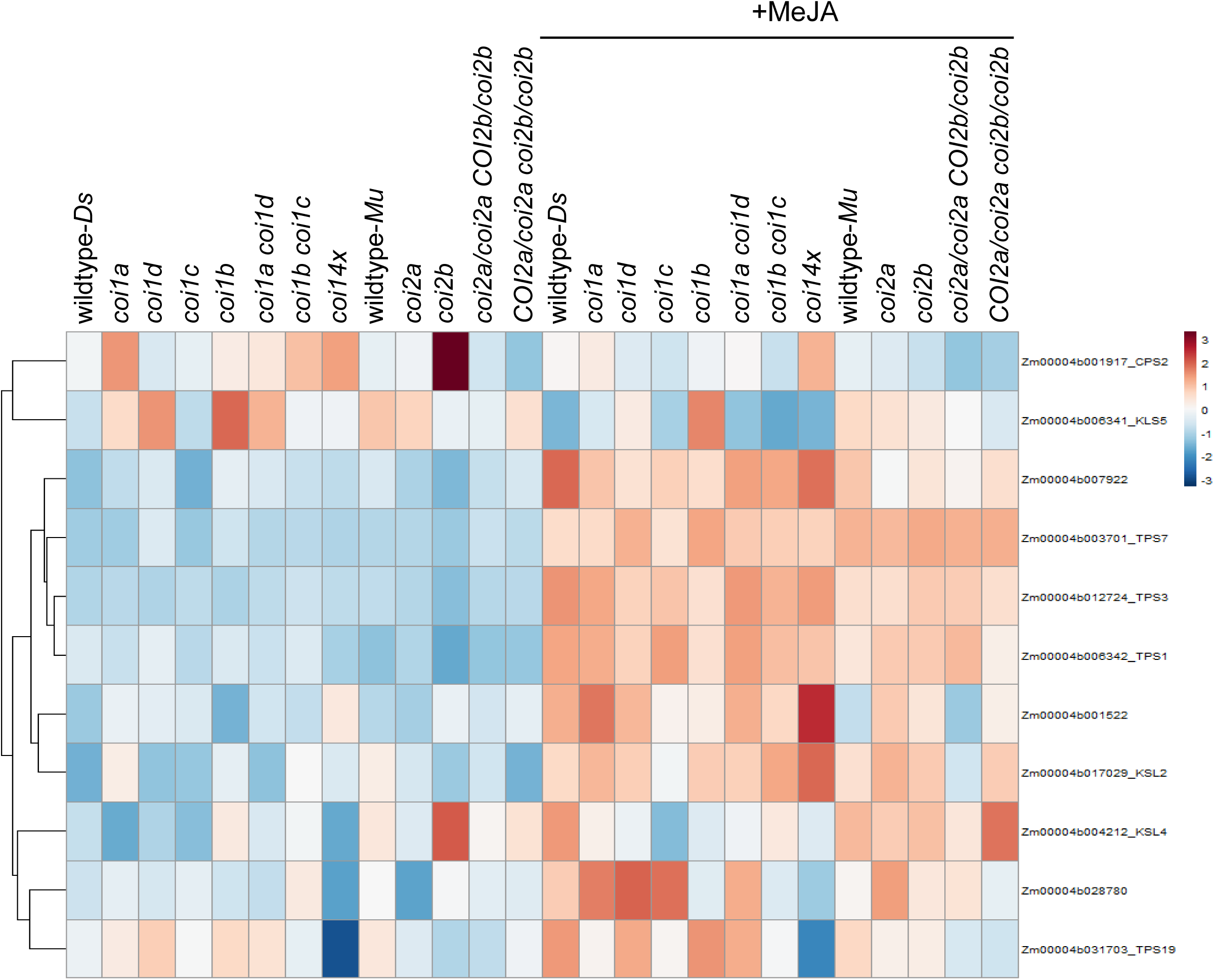
Heatmap showing expression of the terpene synthase genes with and without MeJA induction. The expression levels of the genes used in this heatmap are presented in Table S8.

**Figure S14.**
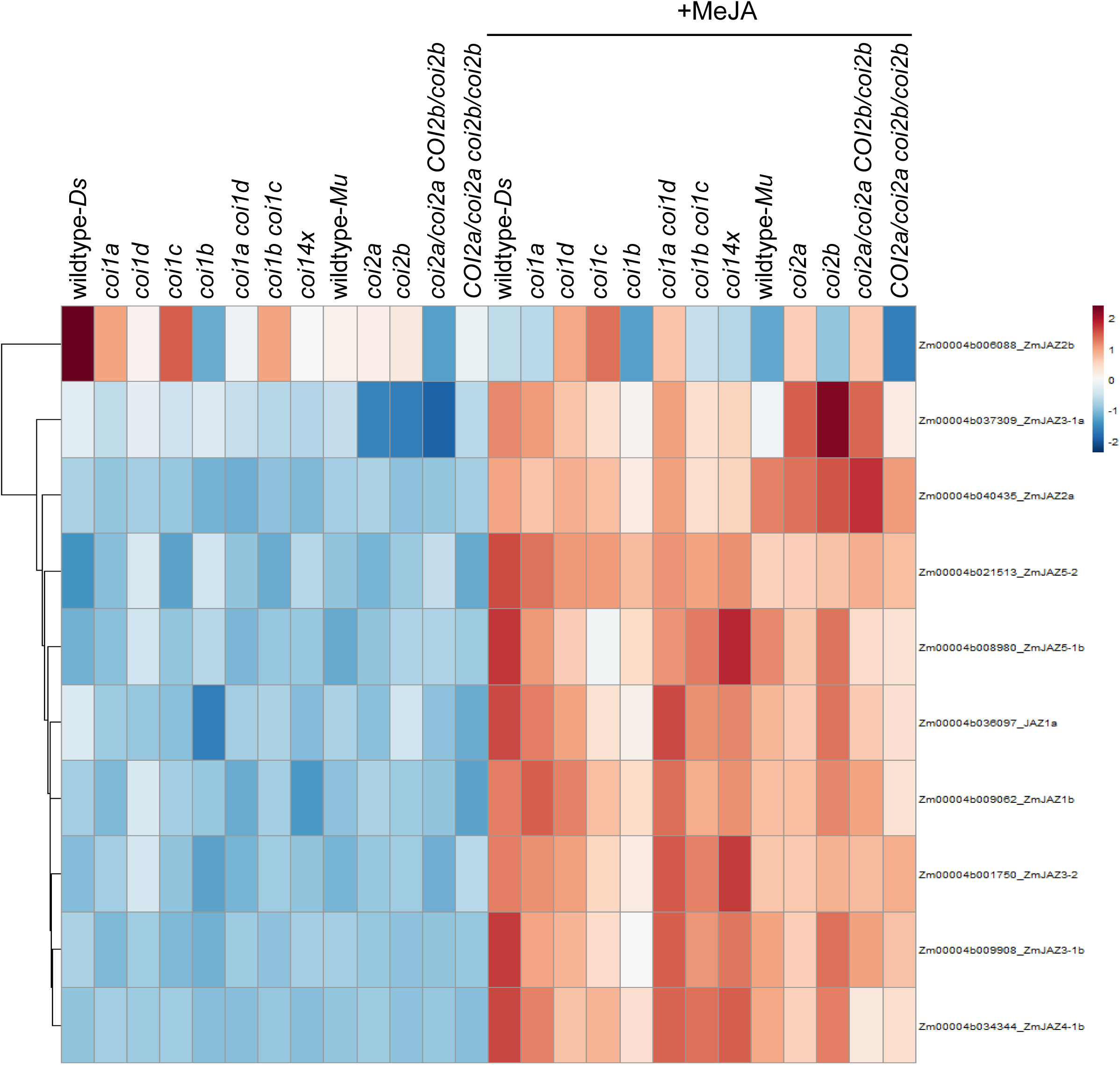
Heatmap showing expression of the maize *JAZ* genes with and without MeJA induction.The expression levels of the genes used in this heatmap are presented in Table S9.

**Figure S15.**
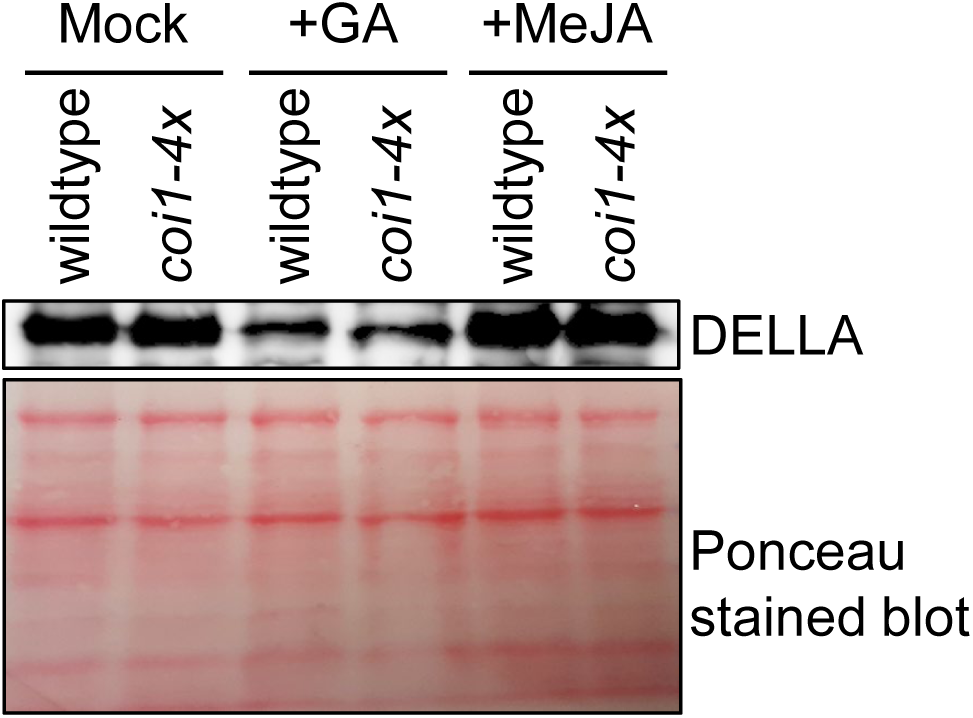
Immunoblot analyses of the maize DELLA proteins after treatment of the 5-day-old seedlings germinated in CYG germination pouches, with gibberellic acid (GA) and methyl jasmonate (MeJA) for two days. The leaf total proteins from equal surface area (similar weight) of *coi1-4x* and wildtype inbred line W22 leaves were analyzed by probing with the rice SLR1 antibody. The membrane was stained with Ponceau S as the loading control.

**Figure S16.**
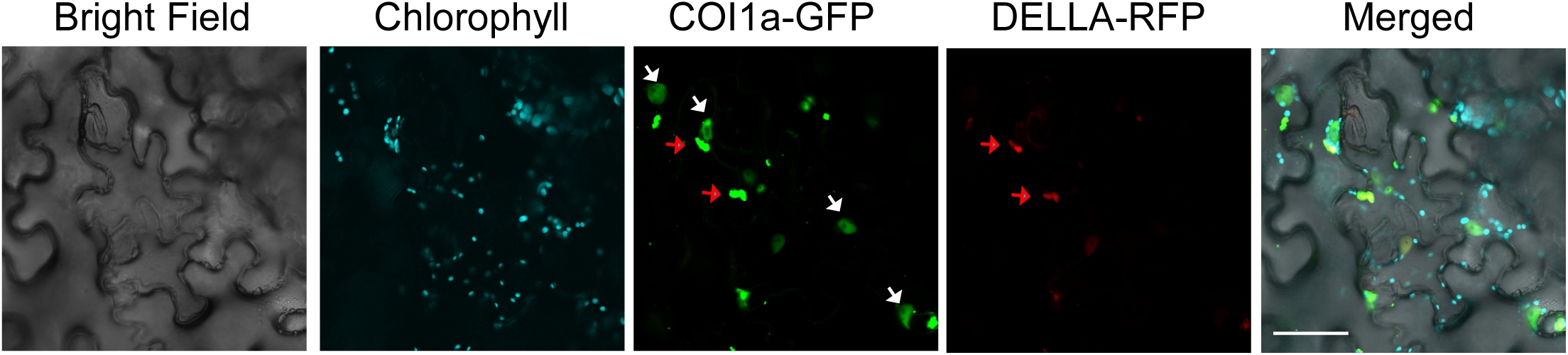
DELLA-RFP colocalizes with the COI1a-EGFP in cytosolic condensates. Whereas the co-expression of the DELLA-RFP with COI1a-EGFP leads to the disappearance of nuclear-localized DELLA-RFP (Figure 11), the cytosol-localized DELLA-RFP is detectable. White and red arrows represent the nuclei and the cytosolic condensates, respectively. Scale bar = 50 μm.

